# Histology-Guided Single-Cell Mass Spectrometry Imaging using Integrated Bright-field and Fluorescence Microscopy

**DOI:** 10.1101/2024.12.03.626022

**Authors:** Alexander Potthoff, Marcel Niehaus, Sebastian Bessler, Jan Schwenzfeier, Emily Hoffmann, Oliver Soehnlein, Jens Höhndorf, Klaus Dreisewerd, Jens Soltwisch

## Abstract

The rapidly evolving field of spatial biology revolves around the analysis of cells in their native microenvironment. This analysis can include morphological features, the presence of specific antigens or gene expression. To add another layer of information, recent methodological advances in matrix-assisted laser desorption ionization mass spectrometry imaging (MALDI-MSI) now enable the untargeted analysis of lipids and metabolites at subcellular resolution. The integration of MALDI-MSI at the single-cell level with established optical modalities, however, relies on an accurate yet intricate co-registration. Here, we describe the integration of bright-field and fluorescence microscopy into a prototype ion source of a state-of-the art MALDI-MSI instrument to obtain lipid and fluorescence microscopy-derived information from the same specimen, hence with intrinsic spatial correlation. We demonstrate the potential of the combined mass spectrometric and optical single-cell analysis on three examples. This includes the visualization of intracellular lipid distributions in macrophages, the introduction of pre-MALDI immunofluorescence staining on the example of murine cerebellum, and the heterogeneity of lipid profiles of tumor infiltrating neutrophils correlated to their individual microenvironments. Overall, the achieved tight correlation of single-cell lipid profiles with morphologic features and protein expression patterns constitutes a powerful resource for cell biology.

## Introduction

Two cells in a tissue or a cell culture are never exactly identical even if they share the same genetic code and microenvironment. Heterogeneity of a cell type can be a consequence of differences in age, ontogeny, or tissue imprinting and result in alterations of morphology, gene expression, molecular composition, and ultimately function. In living systems, individual cells are part of interdependent multicellular structures, i.e. tissues, and undergo a constant exchange within local regulatory networks as part of the almost inestimable complexity that is life. Thus, to characterize the identity and state of a cell, the description should follow a holistic approach and include information from a wide range of analytical techniques ^1^. In recent years, single-cell RNAseq and spectral flow cytometry have emerged as valuable tools allowing to profile cellular heterogeneity within multicellular tissues. Similarly, advances in single-cell proteomics provide information on the heterogeneity on the protein level, albeit with much lower throughput ^2,3^. All of these techniques, however, are destructive and lead to artefactual analyses and, most importantly, to loss of information on the morphology, anatomical, location and spatial relationship of cells. Under the umbrella of spatial biology, the emergence of spatial omics technologies including multiplexed immunohistochemistry and spatial RNAseq allows to overcome some of these obstacles ^4,5^. Yet, these techniques are either highly targeted or limited in their spatial resolving power. A complementary addition to this suite of techniques is the analysis of tissue specimens by high spatial-resolution matrix-assisted laser desorption/ionization mass spectrometry imaging (MALDI-MSI). Herein, finely focused laser pulses are applied to pre-defined pixel positions on the sample surface resulting in spatially defined mass spectra that can be processed to reveal signal intensity maps for hundreds to thousands of analytes in an untargeted manner. Depending on sample preparation, this can include lipids, metabolites, glyccans, or peptides. Recent technical improvements like MALDI-2 and transmission-mode irradiation allowed to decrease pixel size to values of 1x1 µm² and below whilst increasing sensitivity, and thus opened the door to MALDI-MSI analysis at (sub-)cellular resolution ^6–11^. First examples of this single-cell mass spectrometry reveal a strong molecular heterogeneity for lipids and metabolites within clonal populations of cells and seemingly homogeneous tissue regions ^12–15^. In order to contextualize these metabolic phenotypes with existing knowledge on the single-cell level, MALDI-MSI has to be connected to prevalent microscopy techniques. To date, the general application of this contextualization is strongly limited by two main factors: spatial resolving power of MALDI-MSI and a precise co-registration with optical modalities. Especially in the context of tightly packed cellular systems such as tissue, shortcomings in either of these factors lead to erroneous assignment of mass spectrometric information to individual cells and hamper the generation of unambiguous cell specific molecular information ^12,13,16^.

To tackle these issues we present hard- and software developments on the ion source of a commonly deployed MALDI-MSI platform (Bruker timsTOF fleX MALDI-2). Utilizing the free space behind the sample, a transmission mode (t-MALDI) geometry allows to employ a high NA objective for laser irradiation as well as microscopy inside the ion source ^11^. In particular, this enables a direct coupling of bright field microscopy (BF) and fluorescence microscopy (FM) with MALDI-MSI of lipids and metabolites in at (sub-)cellular resolution on a single sample (figure 1A-C). Carefully developed sample preparation protocols (figure 1D) compatible with both modalities allow for the use of small molecule fluorophores as well as IF prior to t-MALDI-MSI analysis. Schematically outlined in figure 1 A-C, the unique design of our ion source utilizes the same optics and sample stage for BF and FM as well as MALDI-MSI, inherently co-registering all modalities with sub-micron fidelity. This unique inherent coupling enables a histology-guided selection of measurement regions of interest (ROI) down to the level of a single-cell and provides a vital link that ensures precise co-registration to other optical modalities. Together with state-of-the-art cell segmentation algorithms, the method provides access to combined morphometric and molecular phenotypes of hundreds to thousands of cells from culture and tissue in a semi-automated fashion.

**Figure 1:**
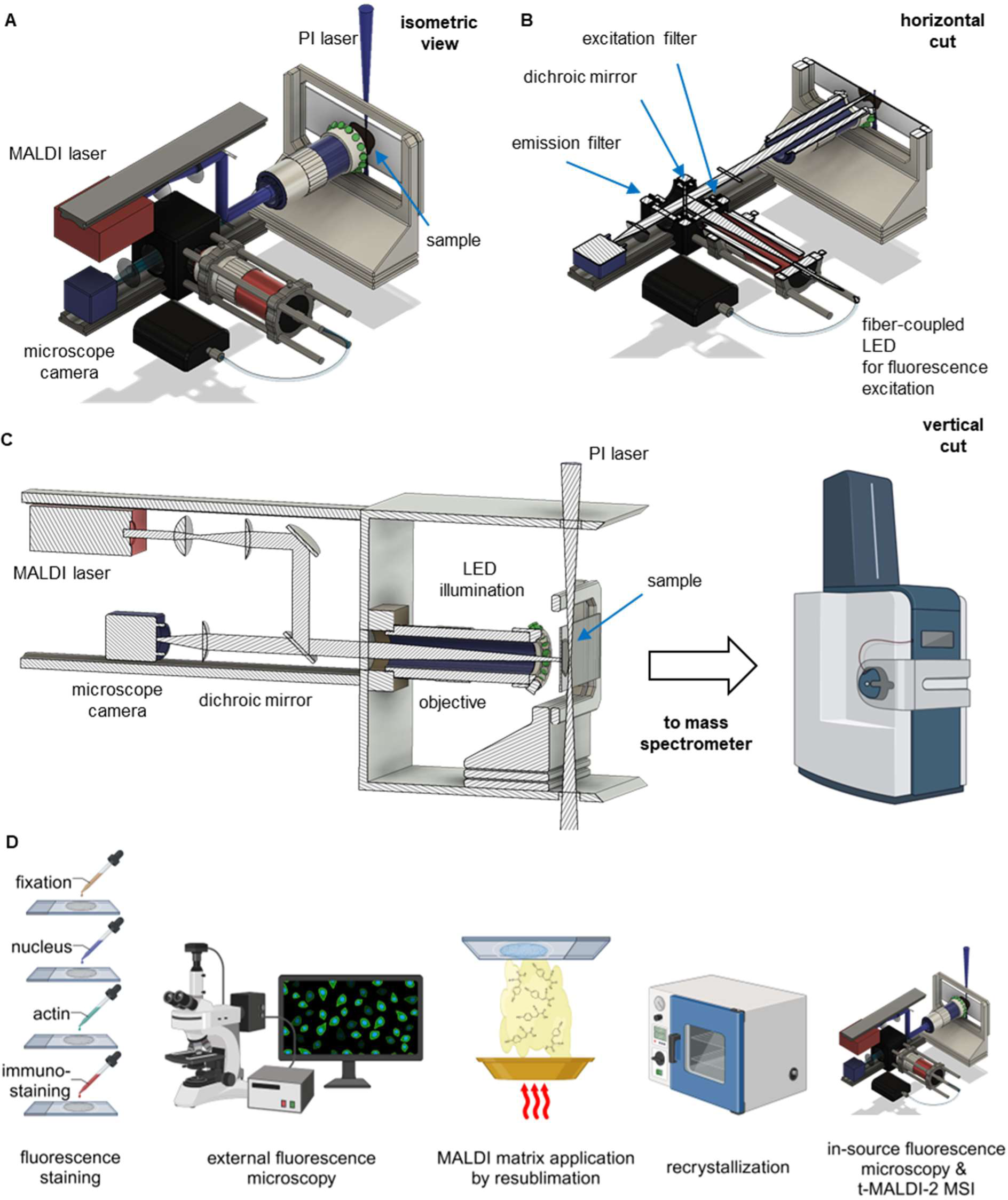
Transmission-MALDI-2 MSI with integrated scanning microscopy. (A) Isometric view of a schematic drawing of the t-MALDI-2 setup illustrating the general layout of the setup. (B) A horizontal cut through the schematic displays the fluorescence microscope components of the setup. Fluorescence is excited by a fiber-coupled LED and filtered by a filter cube consisting of an excitation filter, a dichroic mirror and an emission filter. (C) With a horizontal cut through the schematic the coaligned beam paths of the MALDI laser and the camera are elaborated. The sample is illuminated homogeneously by an LED ring, light from the sample is collected by the objective and focused onto the sensor of a digital camera. The beam of the MALDI laser is coaligned to the first mentioned beam path with a dichroic mirror and is then focused onto the sample by the same objective. Ions generated by the MALDI process with optional MALDI-2 postionisation (PI) are extracted to the mass spectrometer. (D) To enable fluorescence microscopy and t-MALDI-2 on the same sample, different staining steps are performed following a brief fixation. Samples are imaged with an external slide scanning fluorescence microscope, to acquire images of high quality and unperturbed by the MALDI matrix., The samples are then prepared for MALDI MSI by homogeneously coating them with MALDI matrix (Figures S1A, S1B and S1D), and performing a recrystallization step. To enable high fidelity co-registration of the fluorescence microscopy image, a second fluorescence image is taken in the ion source prior to the t-MALDI-2-MSI measurements (Figures S2A – S2C).

## Results

### Transmission MALDI-2-MSI with integrated scanning microscopy

The size of eukaryotic cells typically averages between 15 and 50 µm in diameter, displaying a wide variety in shape and form. Consequently, the spatially resolved study of subcellular details in its morphology or the analysis of larger organelles like the nucleus requires pixel sizes of 2x2 µm² or below. Similarly, the analysis of tight cellular assemblies like tissue requires a sufficiently small pixel size to avoid the sampling of two or more cells within a single larger pixel. In this context, undersampling strongly increases ambiguity in the assignment of signal intensity to an individual cell ^12,13^. In MALDI-MSI, small ablation craters can be achieved by employing laser optics of large numerical aperture (NA) in combination with a sufficiently accurate stage movement with adequate precision and repeatability in the range below 100 nm for the whole region of interest ^11,17^. In this work, we employ a technique coined transmission mode (t-)MALDI where the focusing objective is beneficially placed behind the sample mounted on a transparent glass slide ^18^. The newly developed ion source was coupled to a state-of-the-art orthogonal time-of-flight (oTOF) mass analyzer (Bruker timsTOF-fleX MALDI-2).

To increase sensitivity, the t-MALDI setup is equipped with laser postionization (MALDI-2). This enables the acquisition of information rich t-MALDI-2-MSI of lipids on the cellular and subcellular level ^11,19–21^. While previous iterations of t-MALDI ion sources have hinted at the potential to utilize the employed objective for microscopy of the sample inside the ion source ^11,22^, the setup presented here makes full use of this option and now enables the acquisition of contrast-rich slide scanning microscopy images in BF and FM. A full technical description of the optical setup is supplied in the methods. In brief, the laser is delivered to the sample using a dichroic notch filter with a high wavelength cut-off at 380 nm (Figure 1A and B). Utilizing an apochromatic tube lens and digital camera allows for the microscopic observation of the sample in the near UV and visible wavelength range.

For BF in reflective geometry, the sample is illuminated using a custom-built ring light consisting of 15 LEDs (528 nm) placed around the objective (Figure 1C). Fluorescence microscopy is implemented using filter cubes with an in-line illumination scheme that sources different fiber-coupled LEDs as a fluorescence excitation light source and appropriate sets of excitation and emission filters in combination with the respective dichroic mirror (Figure 1B). The employed custom-built closed-loop sample stage is based on piezo actuators and provides an accuracy of ∼40 nm. In combination with software written in-house that controls camera lighting and stage, this allows for the acquisition of slide scanning microscopy images in BF as well as in FM mode. In BF contrast is provided by an interplay between tissue and applied matrix making the resulting image quality strongly sample dependent (see Figure S1C for examples). For FM on the other hand, images contrast is dictated by the fluorescence properties of the tissue itself and is therefore more independent of the applied layer of matrix (see Figures S2A to S2C). Since FM requires the presence of suitable fluorophores inside the sample, staining methods were developed that are compatible with subsequent MALDI-MSI analysis. For this, protocols were optimized to omit or reduce chemical alterations and spatial delocalization of the analyte content of interest during staining. The typical workflow for sample preparation is depicted schematically in Figure 1D. After a brief fixation and the desired staining, samples are washed, dried, and submitted to external slide scanning microscopy. In a second step, MALDI matrix is applied by resublimation, using a custom-built chamber (Figure S1). Optionally, analyte extraction into the matrix layer can be enhanced by a recrystallization step ^23^.

Inside the ion source, optical beam paths and stage movement for laser delivery and in-source microscopy share essential components. In the xy-plane (sample surface), the site of laser ablation is therefore inherently linked to a specific position on the optical microscope image. In z-direction, along the beam axis, camera image and laser focal planes are aligned. Using these inherent links between laser delivery and microscopic images, the built-in microscope serves for a number of purposes once the sample has been introduced to the ion source. In a first step, an autofocusing routine is used to build a focus map and interpolate the surface topography of the sample within the region of interest. This topography is subsequently used to correct for variations in sample height not only during slide scanning microscopy but also during t-MALDI-2-MSI. This is especially important for the employed high NA focusing objective, with a depth of focus of 1.4 µm. Here, the use of an automatically generated sample topography increases data quality and significantly reduces the time to set up a sample for analysis. In a second step, the internal microscope is used to record slide scanning microscopy images for BF and FM. Because of the inherent spatial link to the laser ablation spot, these detailed images can be used to define regions of interest with high precision, avoiding the recording of off-target and undesired pixels. As a result, the user can accurately choose which cells and tissue microenvironments to investigate based on the fluorescence microscopy.

The size of craters produced during material ejection largely depends on the choice of matrix, the type of tissue and the employed laser power and can range from 0.7 to 1.5 µm ^11^. This enables reliable t-MALDI-2-MSI analysis at a pixel size between 1x1 and 2x2 µm². High scanning speed of the employed stage and optimized and synchronized ion transport and orthogonal TOF-MS geometry allow to record more than 20 pixels per second ^21,24^.

In addition to the described advantages used prior to MSI analysis, the in-source optical microscopy images also provide a crucial direct link between the results of MSI analysis and FM recorded on an external microscope. In conventional setups, different sample preparation steps may induce minor changes in tissue morphology, such as shrinkage upon dehydration. In addition, the information in MSI and FM is produced very differently and from different sources. Together, this can make accurate co-registration of different modalities on a cellular or sub-cellular level exceedingly difficult ^25^. High fidelity co-registration between the in-source and external fluorescence image, on the other hand, is straight-forward. With its inherent link to the MSI data, the in-source image therefore provides a crucial intermediate for fast and accurate co-registration and subsequent correlative analysis (Figures S2A to S2C).

### Lipid analysis at sub-cellular resolving power in cultured cells

To demonstrate the potential of histology guided t-MALDI-2-MSI for the analysis of cultured cells, we investigated THP-1 derived macrophages (see methods for details). THP-1 cells where cultured and differentiated directly in chamber slides and stained for FM using Hoechst 33342 (nuclei), CellMask™ Green Actin Tracking Stain (actin) and LipidSpot™ 610 (lipid droplets). Slide scanning FM was performed using a VS200 slide scanner (Olympus/Evident) with a 50X objective (air) in the DAPI, FITC and Cy3 channels. Subsequently, samples were coated with CHCA matrix for MALDI-MSI analysis (see Methods for details). Using the external fluorescence image (Figure 2A) for a first reference, a region of interest containing 20 cells of heterogeneous morphology was selected. Insight the ion source, the microscopy setup was used to produce a height map based on autofocusing with the DAPI channel as well as a slide scanning images for the DAPI and FTIC channel for the ROI. Subsequently, the ROI was analyzed by t-MALDI-2-MSI at a pixel size of 1x1 µm² (Figure 2B). As previously described, the THP-1 derived macrophages present a strong intercellular heterogeneity (Figure 2C)^12,15^. Image co-registration between the externally generated FM image (Figure 2A) and MSI data (Figure 2B) was performed by using the internal FM as an intermediate as described above (see also Figure S2A in the supplementary information). The overlay of both modalities reveals a high-fidelity co-registration. Cell borders visualized by the actin stain in FM accurately co-align with those reconstructed from lipid signal intensities in MSI revealing little to no leakage or diffusion of lipid content or other preparation artifacts that may compromise cellular integrity ^19^. On the level of a single macrophage, t-MALDI-2-MSI reveals intracellular differences in lipid distribution (Figure 2D).

**Figure 2:**
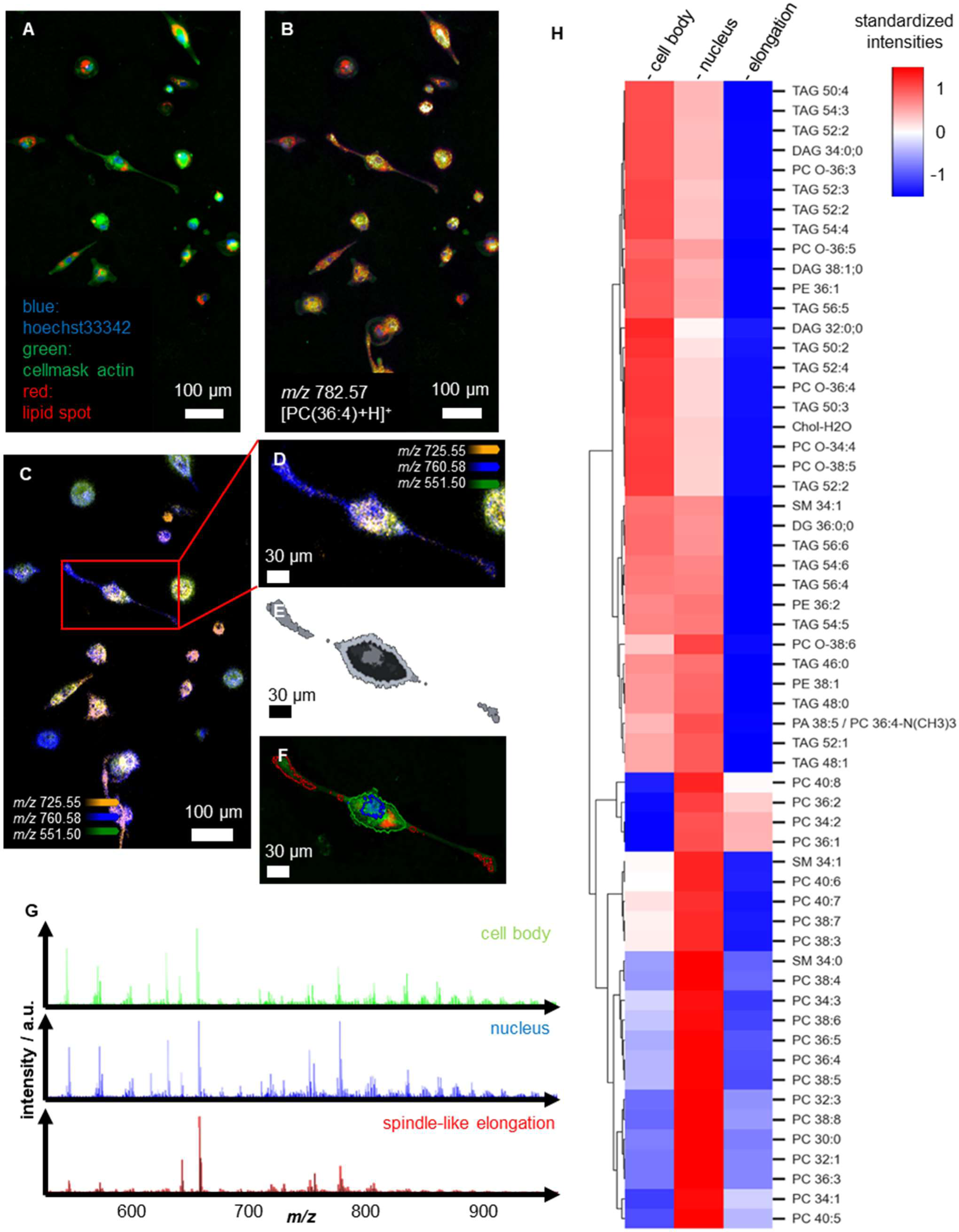
Lipid analysis at sub-cellular resolving power in cultured cells. (A) An external fluorescence image of THP-1 derived macrophages is taken before MALDI matrix application with a slide scanning fluorescence microscope using Hoechst 33342 (nuclei, DAPI channel, blue), CellMask™ Green Actin Tracking Stain (actin, FITC channel, green) and LipidSpot™ 610 (lipid droplets, Cy3 channel, red). (B) After matrix application, t-MALDI-2-MSI is performed and co-registered to the external fluorescence image through the in-source fluorescence image (Figure S1A). As shown on the example of [PC(36:4)+H]^+^ at *m/z* 782.57,the ion signal intensity distribution overlays to a high degree of fidelity with the fluorescence image from (A). (C) An overlay of three different ion signal intensity distributions of [SM(34:1)+H]^+^ (*m/z* 725.55, orange), [PC(34:1)+H]^+^ (*m/z* 760.58, blue), and [DAG(32:0)+H]^+^ (*m/z* 551.50, green) reveals a strong intercellular heterogeneity. (D) A magnification of a small region from the t-MALDI-2-MSI in (C) demonstrates intracellular differences in lipid distribution. (E) With bisecting k-means clustering performed on the MSI data in SCiLS Lab software, distinct clusters could be defined, presenting intercellular differences in lipid ion signal distribution. (F) The regions from the bisecting k-means clustering in (E) co-localize with cellular areas also discernable in fluorescence microscopy. Areas could be assigned to the nucleus (blue), the cell body (green) and the spindle-like elongation (red) by overlaying them with the external fluorescence image from (A). (G) For the regions defined in (F), cell body (green), nucleus (blue), and spindle-like elongation (red), distinguishable mean mass spectra can be extracted. (H) The standardized intensities of different lipid species in the cellular areas is visualized with a heat map.

To reveal cellular compartments with similar lipid signatures, bisecting k-means clustering on the level of single pixels was utilized directly in SCiLS Lab software, revealing distinct clusters. As depicted in Figures 2E and 2F, the resulting clusters co-localize with cellular areas also discernable in FM, namely the nucleus (outlined in blue), the cell body (green) and the spindle-like elongation protruding from the cell (red). Again, the co-alignment of the borders of the clusters identified based on MSI with the outlines of the cellular features visible from FM results confirms the preservation of the cellular structure and integrity during the process of sample preparation. Depicted in Figure 2G, all three clusters that correspond to cellular compartments show distinctly different mass spectra. This is also reflected in a heat map (Figure 2H) that displays the relative distribution for a number of lipid species. Tentative assignments are based on the comparison with results from shotgun lipidomics analysis of cellular extracts from THP-1 derived macrophages ^15^ (Table S1). The heat map reveals that triacylglycerides (TAG) are most commonly detected from the cell body, and, to a lesser extent, from the nucleus region, while they are absent in the spindle-like elongation. Since TAGs have been described to agglomerate in lipid droplets, this observation is in good agreement with the distribution of the lipid spot stain in the FM image (Figure 2F). Some lipid signals are highly increased in the nucleus region; among these are a number of phosphatidylcholines (PC) with polyunsaturated fatty acyl (PUFA) chains. A relatively small number of PCs commonly found in the plasma membrane dominates the elongation region.

### Immunofluorescence-guided spatial lipidomics

To enable a direct alignment of cellular t-MALDI-MSI with more conventional cell typing and identification, a direct connection to fluorescence microscopy on the same sample is pivotal. Here we introduce carefully optimized methods, partially adapted from life cell imaging of cell culture that allow for a MALDI-compatible staining and FM analysis prior to MALDI-MSI (Figure 1D). This includes small molecule stains such as Hoechst 33342 as well as IF staining utilizing either conjugated primary antibodies or systems with primary and secondary antibodies (see Methods for details). To benchmark the newly developed methods, we conducted combined FM and t-MALDI-2-MSI on coronal cryo-sections of mouse cerebellum. This tissue exhibits characteristic histological features across different lateral scales each presenting distinctly different lipid profiles and has been intensely investigated using a wide range of microscopic methods ^26^ and a number of MSI modalities ^27–29^. In addition it is well characterized with regard to its lipidome based on highly sensitive analyzes of lipid extracts ^30^. Figure 3A presents a slide scanning FM image of cell nuclei (Hoechst 33342, DAPI channel), F-actin (CellMask™ Green Actin Tracking Stain, FITC channel) and IF staining of calbindin (Cy3 channel) recorded prior to matrix application in an external slide scanning microscope (VS200, Olympus). Figures 3B and C show the t-MALDI-2-MSI results for a signal assigned as [PE(40:6)+H]^+^ at *m/z* 792.55 for the whole investigated tissue section and a zoomed in region. Uniquely utilizing the internal fluorescence microscopy image enabled a straightforward high-fidelity co-registration of both modalities (see Figure S2B). A partial overlay of MALDI-MSI with the fluorescence microscopy images (Figure 3C) reveals a good correlation with histological features of the white matter and granular layer discernable based on the F-actin staining and the nuclei. Similarly, Figures 3D and 3E show the signal intensity distributions for a signal at *m/z* 834.60 assigned as [PC(40:6)+H]^+^. This lipid has been reported to be strongly expressed in the cell bodies of the Purkinje cell layer ^31^. Here, a comparison with the IF based specific staining of a single Purkinje cell body (Figure 3E) confirms a high fidelity in co-registration between the two modalities and reveals little to no de-localization of the investigated lipid induced during staining or sample preparation.

**Figure 3:**
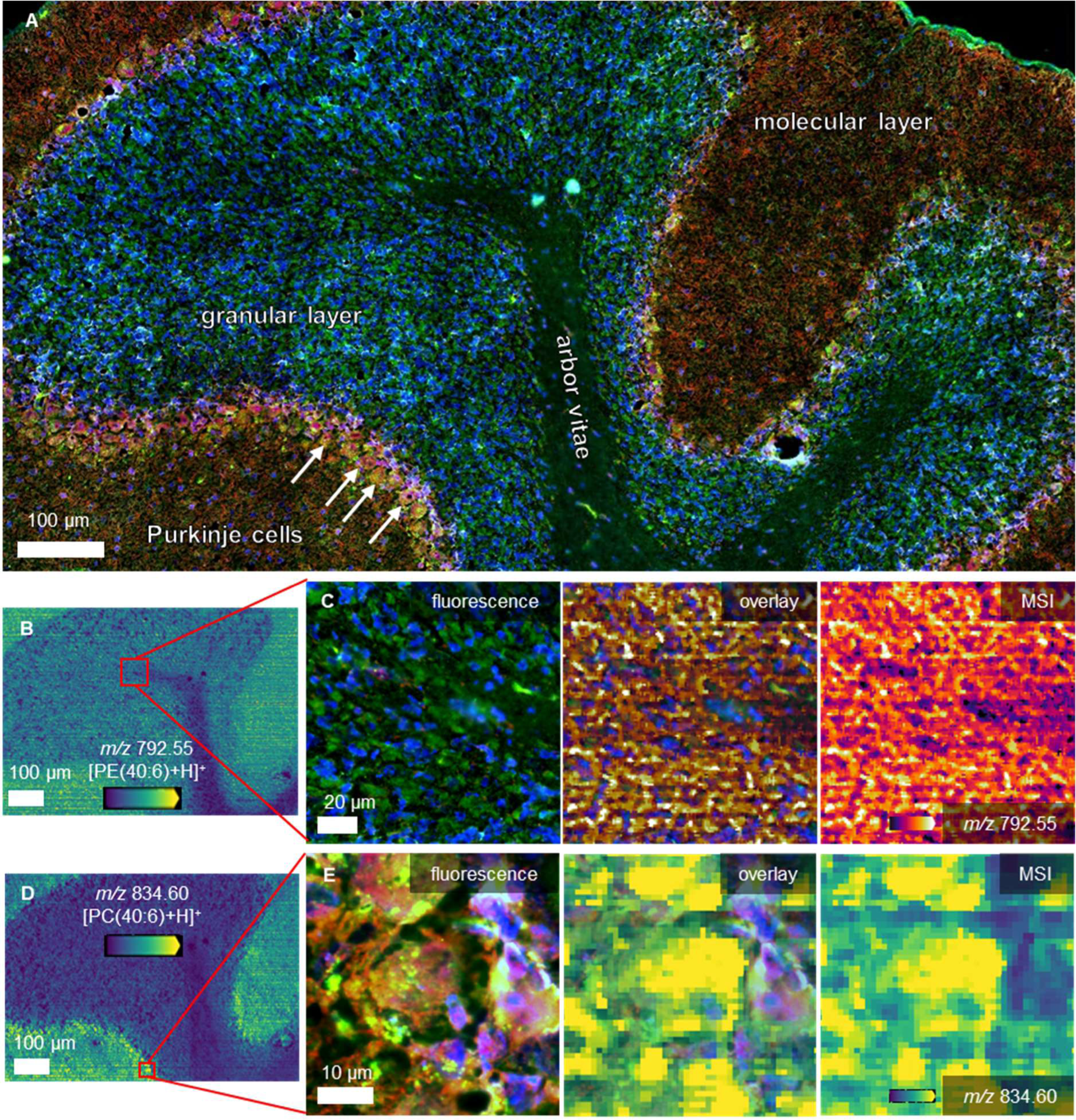
Immunofluorescence-guided spatial lipidomics. (A) An external fluorescence image of an 8 µm section of a mouse cerebellum is taken before MALDI matrix application using Hoechst 33342 (nuclei, DAPI channel, blue), CellMask™ Green Actin Tracking Stain (actin, FITC channel, green) and Alexa Fluor® 594 Anti-calbindin antibody (calbindin, Cy3 channel, red). The different layers of the brain (molecular layer, granular layer, and arbor vitae), as well as the Purkinje cells are well distinguishable. (B) The ion signal intensity distribution of [PE(40:6)+H]^+^ (*m/z* 792.55) measured with t-MALDI-2-MSI shows good contrast over the whole probed region. (C) Overlaying the MSI signal with the fluorescence microscopy image (Figure S2B) reveals a good correlation between the histological features of the white matter and the granular layer. Positions of cell nuclei (blue on the fluorescence image) coincide exactly with the spots of low ion signal intensity in the overlay and MSI. (D) The Purkinje cell layer is found to express a high ion signal intensity of [PC(40:6)+H]^+^ (*m/z* 834.60). (E) The overlay of MSI and FM on a zoom in on a Purkinje cell reveals high fidelity in co-registration and minimal de-localization of the investigated lipid.

To investigate the depth of information for lipid analysis available from pre-stained t-MALDI-2-MSI at a pixel size of 1x1 µm², mass spectral information was matched against a full lipidomics profile for murine cerebellum available from the literature ^30^. As is common for MALDI-MSI, tentative assignment was based on accurate mass and was carried out with a tolerance of 3 ppm. Consequently, this assignment includes the lipid class, the number of carbons and number of double bonds in the fatty acyl chain as well as possible oxidations. It does not, however, differentiate lipid isomers of the same or different lipid classes (see Methods for details).

On this level of specificity, the conflation of isomeric structures deductible from literature results in 255 unique *m/z*-values. Of these, 67 *m/z*-values were also detected in t-MALDI-2-MSI and produced meaningful intensity distributions (Table S2). In particular this includes lipid classes like PE and PS that may be affected for longer formalin fixation times ^32^. Comparison of the detected signal intensities with the molar content of the different lipoforms of each of the detected lipid classes shows a good agreement with the literature (Figures S3A to S3E) and reveals that 89% of the reported molecular lipid content of cerebellum are detected in the t-MALDI-2 measurement ^30^. Lipid species that remain undetected in MSI naturally occur at low to very low concentrations in cerebellum. More pronounced deviations from the literature, such as PC 32:2 that shows a much lower content in MSI may be explained by differences in the employed mouse model; here, the literature is also at variance ^33^. Overrepresentation of other lipid species in the MSI data such as PE 40:7 and PS 40:7 may be explained by the selection of the investigated area or by specific fragments that are produced during the MALDI process, which is most likely the case for diacylgycerols (DAGs). MSI only probes a small region, and site-specific lipids could be missed depending on the measured region, whereas bulk analysis most commonly averages over a much larger volume. Some lipid classes are not detected in the presented t-MALDI-2-MSI data. While some of them occur at very low concentration, such as cardiolipins (CL) or ceramides (Cer), others such as phosphatidylinositol (PI) are commonly not detected in the positive ion mode, which was exclusively used in this study ^34^.

To identify tissue areas with similar lipid profiles, bisecting k-means clustering on the pixel level was performed in SCiLS lab software (Bruker) using the list of *m/z*-values annotated to lipid species discussed above. Figure 4A presents the spatial distribution of the resulting nine most discriminating clusters. A comparison of these areas with histological features based on precise co-registration of the external FM image allows for their classification. Starting with four histologically discernable areas corresponding to the molecular layer (ML), the granular layer (GL) the arbor vitae of the white matter (WM) as well as the Purkinje cell bodies (Figure 4B), either single cluster or a combination of clusters were assigned to each tissue region. Facilitated by the high resolving power of the employed MSI measurement, the outlines of the resulting areas differentiate the Purkinje cell bodies from the surrounding tissue with high accuracy. It may be speculated that the smaller areas of the same lipid profile that surround the cell bodies can be assigned to the complex system of dendrites extending from the Purkinje cells into the ML.

**Figure 4:**
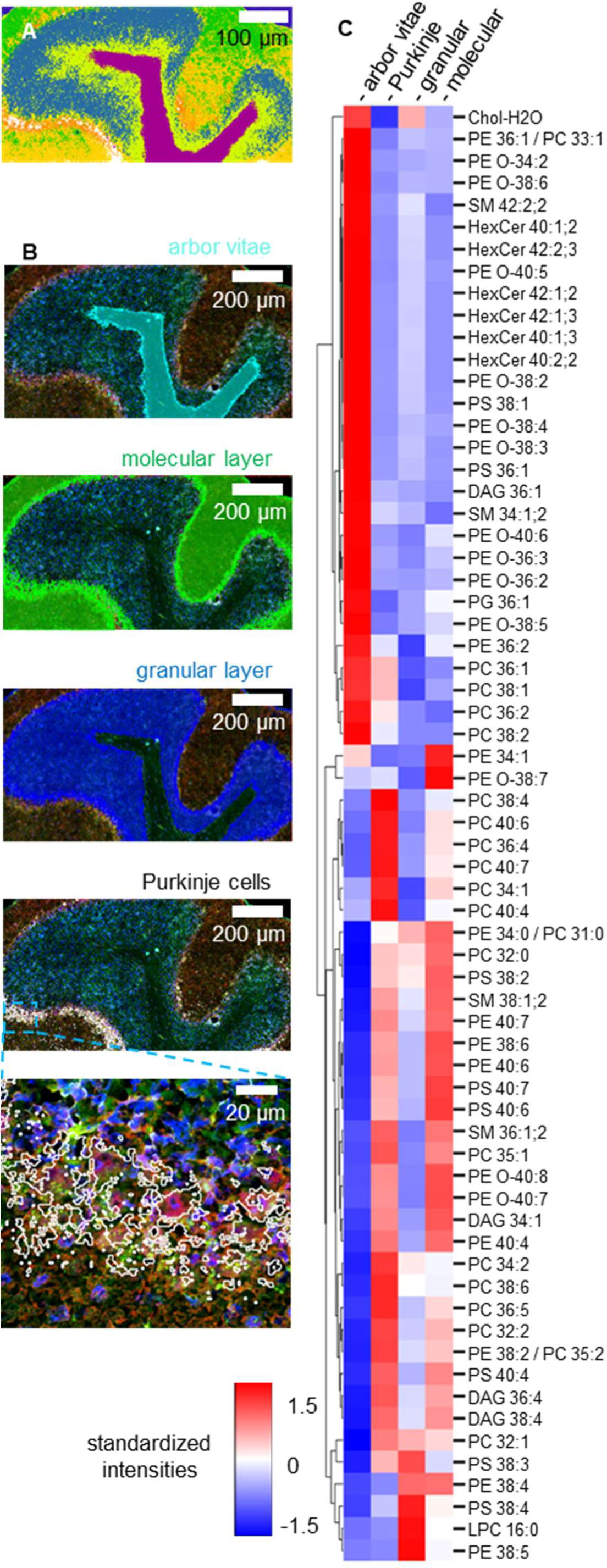
Lipid profile of Purkinje cells and cerebellum layers. (A) By bisecting k-means clustering in SCiLS lab software, 9 discriminating clusters could be created based on the lipid profile from the t-MALDI-2-MSI shown in Figure 3. (B) Clusters from (A) were selectively combined in a way to correlate with the Purkinje cells and cerebellum layers identified by FM. Overlay images of the regions created in that way and the FM image (Figure 3A) are shown for the arbor vitae (teal), the molecular layer (green), the granular layer (blue), and the Purkinje cells (white). For the Purkinje cells, an additional zoom-in is provided to present the excellent correlation of the MSI-based clustering with the red channel from FM. (C) The standardized intensities of the ion signals for 67 lipids are represented as a heat map for the regions identified in (B).

Overall, the high level of discrimination based on MSI overlayed with the respective FM images enables an accurate assignment of lipid profiles to specific cell types or microenvironments. Figure 4C visualizes the lipid distribution of the four tissue types described above in the form of a heatmap. While a more in-depth analysis of the lipid profile of the identified clusters is beyond the scope of this paper, some attributes of the lipid distribution are directly obvious. For example, and in agreement with the literature, hexosylceramides (HexCer) and sphingomyelins (SM) are almost exclusively detected from the WM ^33^. For the cluster assigned to the Purkinje cell layer, a number of phosphatidylcholines (PC) with long chain PUFAS are highly overexpressed as compared to the surrounding tissue ^31^. In contrast, most of the respective PUFAS containing phosphatidylethanolamines (PE) and phosphatidylserines (PS) are mainly detected from the ML and, to a lesser extent, from the GL ^11^.

The presented example demonstrates how information available from bulk lipidomics analysis can be augmented with spatial information. While cluster analysis based on MALDI-MSI data attributes lipid profiles to specific regions within the tissue, information available through FM is used to assign these areas histologically. Notably, the high resolving power available through t-MALDI-2-MSI and high-fidelity co-registration to FM images of the same tissue section allowed for the allocation of lipid profiles down to the level of single cells.

### Automated single-cell analysis of lipids in their histological tissue context

Tumor tissue often contains a large number of different cell types in a wide range of states. It is notorious to create a wide variety of metabolic microenvironments translating to a highly heterogeneous lipid profile across the tissue ^35^. While some of these cells cluster to larger agglomerates of similar molecular profile, others, like infiltrating immune cells, may be found as single, solitary cells spread throughout the tumor tissue. The site-specific molecular profile of these cells, such as neutrophils, is of great importance for their fate and function and is believed to be in constant interdependence with its histological niche or tumor microenvironment ^36–38^. Consequently, the investigation of this “molecular crosstalk” not only requires a molecular analysis at the cellular level but also as a tight connection to histological information about each infiltrating cell as well as its individual microenvironment ^39^. To capture the different cellular states of the investigated cell type in statistically significant numbers, manual annotation on the single-cell level are not feasible and automated analysis becomes essential ^16^.

Here, we used the 4T1 mouse mammary carcinoma model to demonstrate an automated analysis of the lipid profile of a large number of individual neutrophils embedded with histological and molecular information of their respective microenvironment. This tumor model is highly malignant with locally aggressive growth and exhibits metastasis to regional lymph nodes and distant organs, particularly lung, liver, and bone ^40,41^. It has a diverse inflammatory tumor microenvironment with a large number of infiltrating immune cells such as T-cells, macrophages, neutrophils and natural killer cells ^42,43^. Figure 5A shows an overview of a tumor section that was stained using H&E and expertly annotated to identify broader tumor microenvironments.

**Figure 5:**
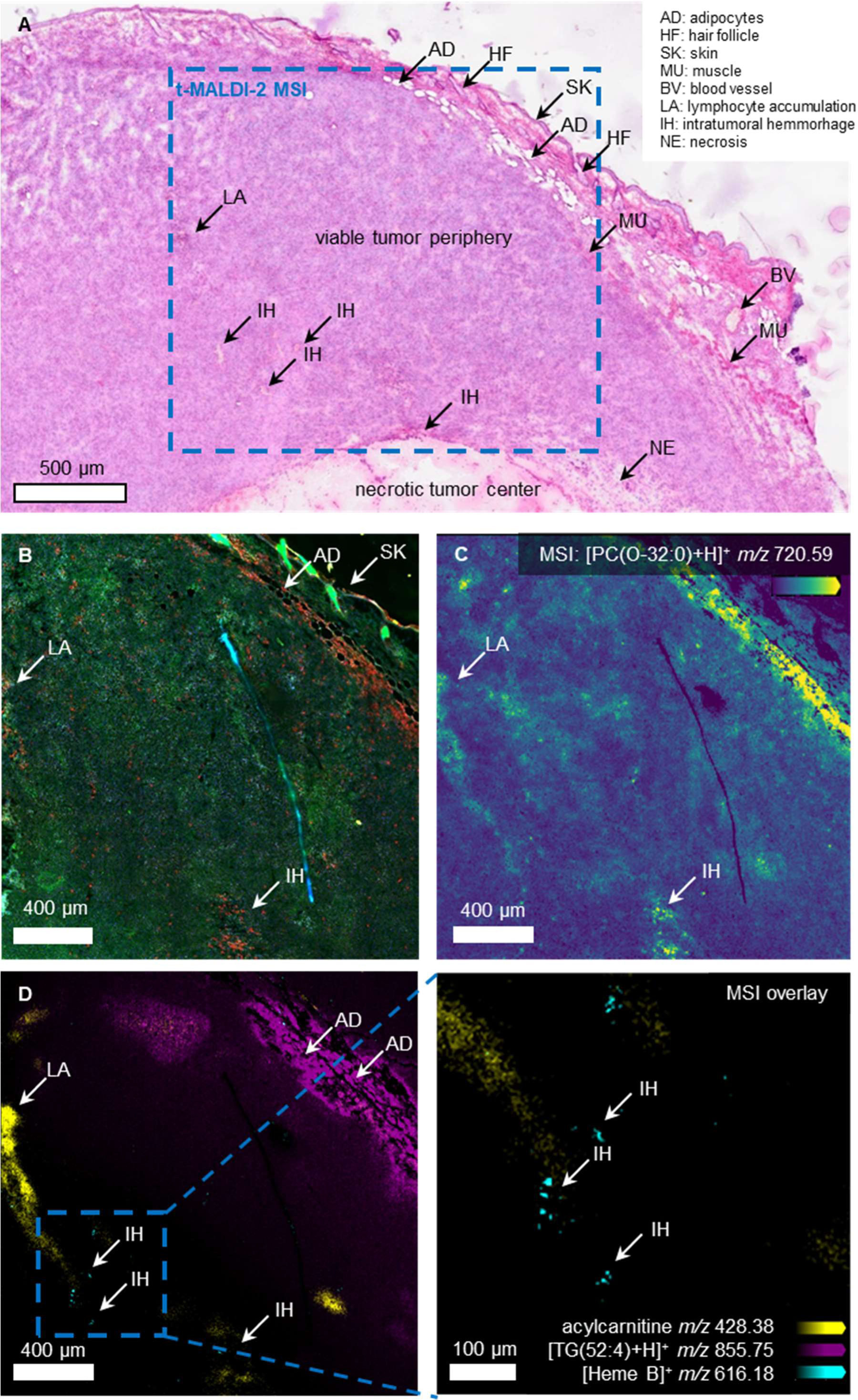
Automated single-cell analysis of lipids in their histological tissue context. (A) To gain an overview of the 4T1 murine tumor tissue, a brightfield image of an H&E stain of a tumor section was recorded and allowed for expert annotation of tumor regions and identification of broader tumor microenvironments. The t-MALDI-2-MSI was conducted on a consecutive section. The region closest to the t-MALDI-2-MSI of the H&E-stained section is marked with a blue rectangle. (B) The section was first stained for FM using Hoechst 33342 (nuclei, DAPI channel, blue), CellMask™ Green Actin Tracking Stain (actin, FITC channel, green) and an IF stain for neutrophils (Anti-Ly6G, Cy3 channel, red) prior to MALDI matrix application and imaged on a slide scanning fluorescence microscope. Based on this image, basic localization of features annotated in the H&E-stained section was transferred. (C) From the t-MALDI-2-MSI, the ion signal intensity distribution of [PC(O-32:0)+H]^+^ (*m/z* 720.59), a marker for neutrophils in mouse, is found to mostly co-localize with the specific Ly6G staining. (D) More ion signal intensity distributions from the t-MALDI-2-MSI from (C), acylcarnitine (*m/z* 428.38, yellow), [TG(52:4)+H]^+^ (*m/z* 855.75, magenta), and [Heme B]^+^ (*m/z* 616.18, cyan), corroborate the histological annotations from (A). Acylcarnitine is a reported marker for hypoxia, TG correlates with the presence of adipocytes, and Heme B can be used as a marker for hemorrhage. This is highlighted in a zoom-in of a region annotated with intratumoral hemorrhage. A selection of additional annotated ion signal intensity distributions can be found in Figure S4 in the supplementary information.

An adjacent section was prepared using MALDI-compatible staining for FM targeting cell nuclei (Hoechst 33342, DAPI channel), F-actin (CellMask™ Green Actin Tracking Stain, FITC channel) and an IF stain for neutrophils (Anti-Ly6G, Cy3 channel) (see Methods for details). All channels were recorded using external as well as in-source slide scanning FM (Figures 5B and S2C) before and after matrix application, respectively. Subsequent t-MALDI-2-MSI reveals the signal intensity distributions for [PC(O-32:0)+H]^+^ (Figure 5C) [acylcarnitine+H]^+^, [Heme B]^+^ and the triglyceride [TG(52:5)+H]^+^ (Figure 5D). The latter three molecular signals can be used to corroborate histological annotations. As expected, strong signal intensity for the observed TG correlates with the presence of adipocytes (compare Figure 5A) ^44^. Acylcarnitine is an important intermediate in lipid metabolism and reported marker for hypoxia ^45^; its signal intensity distribution correlates with an accumulation of lymphocytes detected in the H&E image. Heme B is directly associated with blood and can be used as a marker for hemorrhage ^46^. In addition, spatial distributions for several glycero- and glycerophospholipids reveal a strong heterogeneity across the tumor sample (Figure S4). For one of these lipids, namely PC O-32:0 bulk analysis has identified a sizeable overexpression in neutrophils of the mouse ^47^. In the MSI results presented in Figure 5C, its signal intensity distribution correlates with the specific staining for this type of infiltrating immune cells generally corroborating these findings *in situ*. A direct comparison of MSI and FM on the level of the single cell, however, reveals a heterogeneous expression of this specific lipid throughout the neutrophil population. Figures 6A and 6C show a FM image of a zoomed in region containing three neutrophils as identified by their Ly6G staining (red) and the respective signal intensity distribution for [PC(O-32:0)+H]^+^. Here only two of the three depicted neutrophils also display a strong signal intensity for the lipid species.

**Figure 6:**
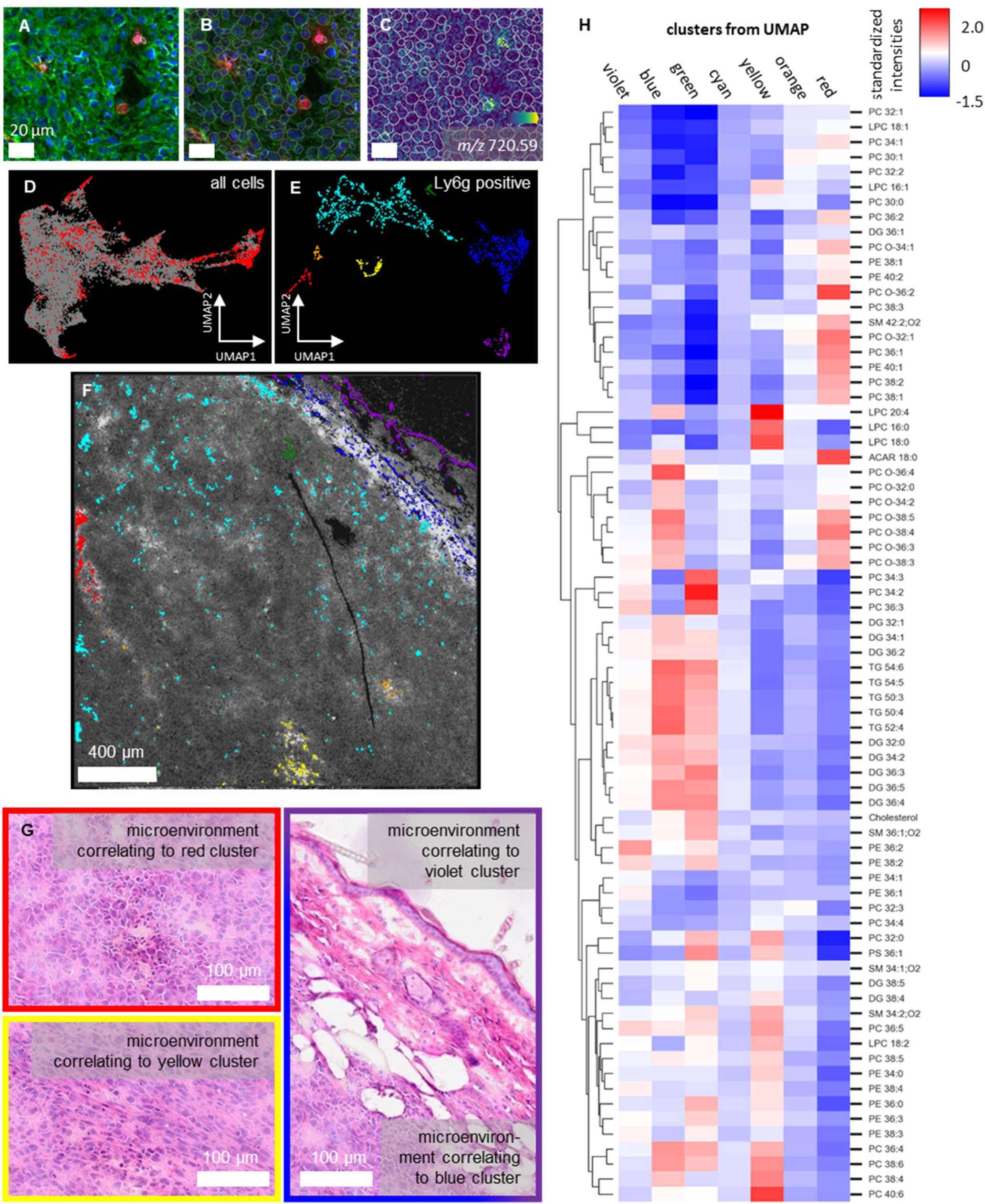
Neutrophil lipid profile in dependence of their state and localization in the tumor tissue. (A) A zoom into a region of the FM image from Figure 5B shows a region where three neutrophils can be determined by the positive Ly6G staining. (B) With DeepCell Mesmer, the contour of 23749 cells (cell masks) are generated on the whole t-MALDI-2-MSI region. Here, only the zoomed in region of (A) is depicted. (C) The cell masks found in (B) are overlayed with the t-MALDI-2-MSI data, here the ion signal intensity of [PC(O-32:0)+H]^+^ at *m/z* 720.59 is exemplarily depicted. Using FISCAS, mass spectra for each cell mask are calculated by combining spectral data of co-localizing MSI pixels. (D) All segmented cells are plotted in a UMAP based on a list of 73 annotated ion signals. Each dot represents an individual cell while the distance between dots visualizes the similarity of the standardized lipid profiles of the cells. Based on the Cy3 fluorescence channel intensity, 1780 Ly6G positive cells were marked in red, while the Ly6G negative cells are presented in grey. (E) All 1780 Ly6G positive cells were filtered and plotted exclusively in a separate UMAP, again using the list of 73 annotated ion species. The UMAP reveals at least seven clearly discernable clusters which are color coded. (F) Cell masks for Ly6G positive cells are projected on a greyscale FM image of the t-MALDI-2-MSI measurement region. The color of cell masks correlates with the clusters found in (E). The projected colored cell masks are not located randomly and can be connected to histological features annotated in Figure 5A. (G) Zoom-ins of regions correlated with the Ly6G positive cell mask clusters from (E) and (F) on the closest corresponding regions on the H&E stained section from Figure 5A allow to connect histopathological context to the different lipid profiles of neutrophil cells. (H) The standardized intensities of 73 ion species for the seven clusters from (E) is presented as a heat map to visualize differences in lipid profiles between the different groups.

To enable the statistical analysis of a general heterogeneity in the lipid profile of neutrophils in combination with their histological context, we adapted an existing workflow described for the analysis from single cells in culture to tissue analysis ^12,15^. For this, the FM image was co-registered to the t-MALDI-2-MSI dataset using a Python adaptation of the Insight Toolkit (ITK) SimpleITK. Taking advantage of the inherent link to the MSI data, the in-source fluorescence image was used as an intermediate step during this registration (Figure S2C). The resulting transformation matrix was applied to resample the FM images to fit precisely the area measured by t-MALDI-2-MSI. These images were submitted to cell segmentation using the Mesmer model from DeepCell ^48^. The segmentation masks were then used to calculate a variety of morphometric parameters using CellProfiler, adding to the wealth of information available for each cell ^15,49^. This data includes parameters describing cell size and shape but also fluorescence intensity for all employed channels. Utilizing the described co-registration, Figure 6B and 6C depicts the outlines generated by the automated cell segmentation projected onto the FM and MS image of a zoomed in area, respectively. The overlay demonstrates the high-fidelity co-registration for the relatively small neutrophils. For subsequent data analysis, a peak list containing 73 entries was used (Table S3). All peaks are tentatively annotated based on exact mass (< 3 ppm). To generate single cell data, signal intensities for the entries of the peak list for all pixels allocated to a specific cell by the respective projected outline are summed (see Methods for details). Pixels touching two or more cells are not included for either cell. For each cell, this data was normalized to the sum of all signal intensities for the respective specific cell. For the whole tissue, cell segmentation and data analysis resulted in combined morphometric and MS data for 23749 individual cells.

Figure 6D depicts a Uniform Manifold Approximation and Projection (UMAP) for all segmented cells. Each dot represents an individual cell and the distance between dots visualizes the similarity of the standardized lipid profiles of the respective cells. To identify neutrophils, the combined data was filtered using Otsu’s method based on FM signal intensity in the Cy3 channel. This has resulted in 1780 cells positive for Ly6G highlighted in red within the UMAP. Interestingly, the neutrophils do not form a separable cluster. This can be interpreted as a high molecular heterogeneity within the neutrophil population induced by a molecular interplay with their respective microenvironments. To investigate this heterogeneity in more detail, Figure 6E depicts a UMAP compiled exclusively for those cells identified as neutrophils. It reveals at least seven clearly discernable clusters. Each of these clusters is marked by a different color and displays a distinctly different lipid profile visualized in the form of a heatmap in Figure 6H. To investigate possible correlations of the molecular profile of these clusters with histological annotations, outlines for all neutrophils where projected onto the FM image of the investigated tumor using the colors assigned to their respective clusters. Neutrophils from each cluster are not randomly distributed across the tissue but can be assigned to specific microenvironments. Neutrophils marked in cyan are found across the viable tumor periphery. Apart from a few smaller agglomerations, they are relatively solitary and most probably represent tumor infiltration. The violet and blue cluster correlate with dermal and adipose tissue (compare Figure 6G) and display an overexpression of TAGs and cholesterol. The green cluster does not correlate with any histological feature visible in the H&E stain, its close proximity to the adipose tissue and the increased levels of TAG in its lipid profile, however, may suggest a subpopulation that recently infiltrated into the tumor from the direction of the adipose tissue. Neutrophils from the red cluster are localized in a hypoxic region that displays an accumulation of lymphocytes (Figure 6G). Neutrophils from this region are comparatively rich in PC-O lipids. The yellow cluster is located near the necrotic core. Histologically, its respective microenvironment displays some lymphocyte accumulation (Figure 6G), less hypoxia as compared to the red cluster, and a relatively high level of detected LPC 20:4 (Figure S4) suggesting inflammation ^50^. Neutrophils associated with this cluster display an increased content of LPC and PUFA containing PC species. The tumor microenvironment of the orange cluster cannot be characterized based on the available information. In future studies, additional IF markers could help to deduce more detailed information on the specific microenvironments. From a molecular point of view, the placement of the orange cluster in the UMAP between the cyan and the red cluster, however, hints towards an intermediate between these two states. All of these observations only provide a first glimpse into the potential of the technique to investigate the interplay between infiltrating immune cells and their respective microenvironments. A more in-depth interpretation regarding the regulation of lipids in neutrophil differentiation and its intricate interplay with the respective tumor microenvironment is, however, beyond the scope of this resource paper.

## Discussion

Modern MS analysis can provide specific molecular information of single cells, thereby delivering an important puzzle piece towards the understanding of processes like cellular differentiation or progression of disease on the cellular level. For a more holistic approach, however, lipidomic or metabolomic information has to be put into the context of the cell type, differentiation status and cell state but also its current tissue macro- and microenvironment ^1^. In this regard, single-cell MS-imaging has been reported to combine the annotation of single cells and their microenvironments *in vitro* provided by (immuno)fluorescence microscopy on the one hand and the analysis of lipids and metabolites at cellular resolving power on the other ^16^. In this paper, we demonstrate that t-MALDI-2-MSI in combination with in-source FM and dedicated sample preparation techniques can lay the groundwork for the large scale, automated, and reliable application of this technique. A pixel size down to 1 X 1 µm², enabled by transmission-mode laser irradiation, permits spatial sampling rates necessary for single-cell resolving power on tissue. The use of laser postionization in combination with sensitive ion optics supply the essential depth of analytical information for a wide range of lipids and metabolites. In-source FM and BF provides conjunct optical analysis. In addition, the resulting inherent link between MS imaging and microscopy, made possible by dedicated MALDI-compatible sample preparation strategies, serves as a critical bridge to other analytical modalities. This tight connection to optical microscopy analysis has long since been identified as a key feature in the integration of MALDI-MSI into the “biomedical toolbox” ^51^. Conventionally, microscopy prior to MALDI-MSI is restricted to the limited analytical capabilities offered by optical microscopy of unstained tissue, e.g. by autofluorescence or phase-contrast microscopy and sample staining is carried out after MALDI-MSI analysis ^52–54^. Application and removal of the matrix and especially laser irradiation that precedes microscopy, however, can damage the tissue, distort images and impede binding properties for antibody based analysis ^8^. High quality microscopy analysis, including IF, is therefore usually carried out on adjacent tissue sections thereby introducing well-known difficulties in accurate co-registration and in the analysis of specifically targeted single cells ^25,51,55,56^.

For the molecular investigations of cell culture models, the technique extends existing single-cell MSI workflows beyond intercellular heterogeneity and towards the analysis of intracellular molecular distributions. In a proof-of-concept study, we demonstrated this feature on THP-1 derived macrophages by discerning lipid profiles for different areas of the cell that correlate with histological features, like the nucleus or the spindle-like elongations. Strong spatial correlation between the two imaging modalities also served as confirmation and quality control for the optimized sample preparation protocols, as very limited delocalization of lipid compounds was discernable. In the future, the technique may be a valuable asset to investigate intracellular metabolic and lipidomic changes in processes like mitosis, apoptosis, or ferroptosis, but also to provide insights into the molecular make-up of relatively large organelles like vacuoles, lipid bodies, or phagosomes.

While single-cell analysis from cell culture grown to sub-confluence has been established to different degrees of automation, work on large-scale single-cell-MSI from tissue sections has been limited ^12–15,57,58^. To some degree, this can be attributed to shortcomings in segmentation of cells based on MSI data alone and/or insufficient co-registration with optical modalities acquired from the same or often even adjacent sections. Consequently, this has forced the manual annotation of cells and limited the respective studies to small cell counts. To disentangle metabolic heterogeneities and to identify rare subpopulations among the same types of cells, however, the analysis of statistically significant numbers is essential ^16^. In addition, the minimal pixel size in commercially available instrumentation lays at 5 X 5 µm². Employed in MALDI-MSI studies for single-cell analysis of tissue, this often leads to a co-sampling of two or more cells in one pixel and results in ambiguities during the compilation of single-cell data.

In the analysis of two models for tissue samples, we have demonstrated how histology guided t-MALDI-2-MSI can help to overcome these shortcomings. In the first step, we investigated mouse cerebellum that is well characterized with regards to its morphology and lipid contents. Comparison with the pertinent literature confirmed that the newly developed sample preparation protocols do neither notably compromise morphological integrity nor the lipid composition of the samples. Analysis of the acquired mass spectra resulted in the annotation of 26% of all previously described lipid species in the murine cerebellum, accounting for about 89% of total molar lipid content. Relative intensities of lipid species within lipid classes are, to a large extent, reproduced correctly. Based on signal intensity distributions of the annotated lipids, we were able to cluster specific areas of the tissue to common molecular profiles and assigned these areas to histological features observed in the FM data with high spatial fidelity. This direct correlation of histological features with signal intensity distributions in MSI served as quality control for the integrity of tissue features down to the level of single Purkinje cell bodies and confirmed the spatial integrity of the investigated tissue for the investigated lipids on the cellular level. Overall, the results demonstrate that dedicated and careful sample preparation enables a subsequent FM and t-MALDI-2-MSI analysis on the cellular level without a sizeable distortion or loss of information from the same tissue section. Notably, we were able include IF into the pre-MALDI staining enabling the specific staining and identification of selectively targeted cells prior to MSI-analysis for the first time.

To demonstrate the analysis of scarce and isolated cells, we engaged in the analysis of tumor tissue generated from a 4T1 mouse model. Here we investigated infiltrating immune cells and their molecular heterogeneity in the context of their individual tumor microenvironment. For this, automated generation of single-cell mass spectra from tissue was achieved utilizing tools for cell segmentation recently introduced for IF microscopy. Exploiting the inherently accurate spatial co-registration between FM and MSI, cell masks provided by cell segmentation were projected onto MSI data, resulting in the generation of single-cell data for a total of 23749 cells from a single MSI measurement. By filtering this wealth of data for Ly6G positive cells, we were able to dissect 1780 neutrophils from the measured region of the tissue sample. Statistical analysis revealed a strong heterogeneity within this class of cells with each sub-population presenting distinctly different lipid profiles. Back-projection of the resulting clusters to their respective spatial origin in combination with histological annotation of the tumor based on microscopy and MSI-data resulted in a histological placement of the different clusters into specific tumor microenvironments.

The seamless integration of morphometric data with single-cell mass spectra significantly expands the informational space available for each individual cell inside the tissue and enables a direct contextualization with its tissue microenvironment *in situ*. In future applications this will allow for the combination of IF based information that may describe the expression of individual proteins or regulation of specific genes^59^ with the expression of lipids or metabolite in the same cell and its direct surrounding. Overall, the presented techniques represent a new and versatile suite of tools for an improved integration of single-cell MSI into existing workflows of modern cell biology. In this, it may, for example, help to decipher heterogeneities within populations of stem cells with regard to cell fate, to define extended molecular phenotypes in the classification of immune cells to predict and understand pro- and anti-inflammatory behavior, and many other possible applications.

### Limitations of this study

Data analysis performed in the scope of the paper is not intended to reveal any new insights into molecular or cellular biology but rather to initiate and trigger more specific work that utilizes the new tools. Molecular analysis presented here is, for the most part, limited to lipidomics analysis. While MALDI-2 MSI is generally not limited to this group of analytes, expansion to e.g. metabolomics or N-glycan analysis, however, will require modifications and adaptation of protocols for sample preparation especially regarding sample diffusion and sensitivity. Inherent to all MS imaging techniques, annotation of molecular species based on the accurate mass alone is limited to a certain degree of specificity and certainty. Complementary analysis of extracts from bulk or on tissue tandem MS experiments are therefore advised to decrease ambiguity in assignment.

While the analysis of single cells from tissue sections allows for a placement into their histological and molecular niche or microenvironment, this placement is limited to the two dimensions of the section. Especially for scarce or solitary cells, however, important features may be found in their three-dimensional vicinity. While 3D MALDI-MSI can be done, it requires strongly increased measurement times and introduces additional challenges in sectioning and co-registration of adjacent sections. By concept, MALDI-MSI is a destructive method that does not allow for live cell imaging or any type of biological activity of cells or tissue after analysis.

## Methods

### Cell culture preparation

THP-1 ACC-16 cells (DSMZ) were cultured in RPMI 1640 Medium (Lonza), which was supplemented with 2 mM L-glutamine, 10% fetal bovine serum and 1 mM sodium pyruvate. About 10 000 cells per chamber were sown in Millicell EZ Slide 8-well chamber slides (Sigma-Aldrich). Differentiation to M0 macrophages was induced by exposure to 20 ng/ml phorbol 12-myristate 13-acetate (PMA) for 24 h followed by a 3-day rest phase in PMA-free medium. M0 macrophages were polarized to M2a macrophages using IL-4 and IL-13 (both Sigma-Aldrich), each 20 ng/ml, for 48 h. Cells were fixed 5 min using 4% formaldehyde. Actin, nuclei and lipid droplets were stained according to the previously described protocol by Bien *et al*. using Hoechst 33342 (Sigma-Aldrich), CellMask™ Green Actin Tracking Stain (Thermo Fisher), and LipidSpot™ 610 (Biotium) ^19^. After staining, cells were washed using 150 mM ammonium acetate and dried at room temperature for 2 h.

### Mouse model

All animal experiments were carried out in accordance with local animal welfare guidelines approved by the responsible authorities (Landesamt für Natur, Umwelt und Verbraucherschutz NRW, Protocol No. 81-02.04.2018.A010). Female BALB/c mice (Charles River Laboratories) were used at the age of 8 to 12 weeks. Mice were housed under a 12-hour light/dark cycle with ad libitum access to food and water. For tumor implantation, 106 4T1 cells were resuspended in 25 µl of DMEM (Thermo Fisher) cell culture medium and then injected orthotopically into the lower left mammary fat pad of the mice. Tumors were allowed to grow for 9 days and tumor size was monitored daily with a digital caliper. On day 9 after tumor implantation, the animals were sacrificed. Brain tissue was harvested from untreated mice used as control group for the mentioned study.

### Tissue sectioning

Mouse brain and tumor tissue were embedded in Epredia™ M-1 Embedding Matrix (Thermo Fisher) and snap frozen in liquid nitrogen. 10 µm (cerebellum) or 8 µm (tumor) thick tissue sections were produced with a cryostat (Leica Biosystems) at -20 °C. Sections were thaw-mounted onto SuperFrost glass slides (Thermo Fisher) for H&E-staining or IntelliSlides (Bruker) for t-MALDI-2-MSI measurements. Samples for t-MALDI-2-MSI were then vacuum-sealed using a food vacuum sealer and stored at -80 °C.

### Tissue fixation and fluorescence staining

Frozen tissue sections were passively brought to room temperature in the vacuum-sealed package. After opening the seal, sections were further dried under a gentle nitrogen stream for 10 minutes. With a Super PAP Pen (Science Services) a contour was drawn around each section, followed by another 10 minutes drying under a gentle nitrogen stream, to create a hydrophobic barrier and thus enabling pipetting aqueous solutions on and off the sample. The following washing, fixation and staining steps were all performed by pipetting ∼100 µl of the accordant solution onto the sample and removing it after the specified time. Slides with tissue sections were placed on a metal plate standing in ice water. Samples were quickly fixed in 4 % formaldehyde in PBS for 5 minutes and washed with PBS for 30 seconds. After tissue fixation, the primary antibody, either Anti-Ly6g or Anti-Calbindin (both rabbit, abcam), diluted 1:500 (v/v) in 1% BSA in PBS was applied to the sample and incubated for 60 minutes. The sample was then washed 3 times for 30 seconds in PBS. In case of a dual antibody staining, the secondary antibody conjugated with Alexa Fluor® 594 (goat anti-rabbit, abcam), diluted 1:1000 (v/v) in 1% BSA in PBS was applied and incubated for 60 minutes, followed by 3 times 30 seconds washing steps in PBS. 5 µg/ml Hoechst33342 (Sigma-Aldrich) and 1X CellMask™ Green Actin Tracking Stain (Thermo Fisher) diluted in PBS were applied and incubated for 5 minutes. After staining, the slide was lifted from the metal plate and parts of the hydrophobic barrier were removed at the bottom of the slide with a pipette tip. Samples were then washed with 2 ml of ammonium acetate each, by letting it flow carefully over the section. For final drying, samples were put under a gentle N_2_ stream for 20 minutes.

### Tissue H&E staining

Hematoxylin and eosin (H&E) stains of tissue sections were performed on untreated consecutive sections directly after sectioning. Sections were stained for 30 s with hematoxylin (Sigma-Aldrich) with subsequent bluing in tap water for 2 minutes. Afterwards, the tissue was stained for 30 s with eosin (Sigma-Aldrich) and then rinsed with water.

### External microscopy

Tissue sections mounted on IntelliSlides (Bruker) and cells mounted on Millicell EZ Slide 8-well chamber slides (Sigma-Aldrich) were scanned using a VS200 slide scanner microscope (Olympus/Evident). H&E stained tissues were recorded in bright field mode using a 20X objective, fluorescence images were recorded at 50X using the DAPI, FITC and Cy3 channels. Image processing was performed with OlyVIA (Evident).

### Transmission optics setup

A schematic of the transmission optics setup can be found in Figure1A-C of the main article. The sample is mounted on a three axis piezo stage (SmarAct) and is illuminated homogeneously by 15 green (528 nm) LEDs (Thorlabs) mounted on top of a 50X objective (Mitutoyo) with a self-designed 3D printed LED ring. Light from the sample is collected by the objective and is focused by a tube lens (Thorlabs) onto the sensor of a digital camera (The Imaging Source). The beam path of the MALDI laser of the timsTOF fleX mass spectrometer is rerouted through a 2:1 telescope optic and co-aligned to the microscopy path with a dichroic mirror (Laseroptik). For recording fluorescence microscopy images, a filter cube (Thorlabs) with a filter set (Thorlabs) for the suitable fluorescence channel (DAPI, FITC, mCherry) consisting of excitation and emission filters and a dichroic mirror is mounted in front of the tube lens. In fluorescence mode, the LED ring is turned off and fluorescence is excited by the respective fiber-coupled LED (Thorlabs) mounted with a collimating optic directly to the filter cube. The camera is integrated in the software by using the IC Imaging Control 4 SDK (The Imaging Source). The trigger signal for image acquisition of the camera and fluorescence excitation is provided as a “position reached” event by the controller of the piezo stage and is handled by a digital delay pulse generator (Quantum Composers). With this pulse generator, the fluorescence excitation time was set from 10 ms to 2 s per image depending on the sample and fluorescence channel.

### MALDI-matrix coating by resublimation

To achieve a reproducible and homogeneous coating of the sample slides with MALDI-matrix, a custom-designed resublimation chamber was developed. A schematic of this setup can be found in Figures S1A and S1B in the supplementary information. Under atmospheric pressure conditions, the hot-side is preheated to 80 °C and 0.5 ml of 10 mg/ml CHCA MALDI-matrix (Sigma-Aldrich) in acetone solution is pipetted into the matrix-reservoir. With the acetone evaporating rapidly, the MALDI-matrix is left forming a uniform layer on the bottom of the matrix reservoir. The sample is then clamped onto the cold-side at room temperature and the whole device is sealed and evacuated. After reaching the final pressure of 10^-2^ mbar, the cold-side is first cooled to -18 °C, then the hot side is heated to 150 °C to start the sublimation and resublimation. After 60 minutes, the heating of the hot-side is turned off and the cold-side is heated carefully to room temperature before venting the system to prevent condensation of ambient humidity on the sample. The sample is then put onto a room temperature metal block in a saturated atmosphere of 0.5 % ethanol in water at 70 °C for 1.5 minutes for recrystallization.

### Autofocus of optics

The MALDI laser and microscope camera utilize the same objective for focusing and observation in the transmission-MALDI setup deployed in this work. A section view schematic of the setup can be found in Figure 1C in the main article. The focal plane of the MALDI-laser is adjusted to match that of the camera by fine adjusting the second telescope lens of the laser beam path. After aligning the focal planes, all optics are kept at a constant position. The objective (Mitutoyo) has a small focal depth of 1.4 µm and samples have to be in focus during optical microscopy and MALDI-MSI. This can be achieved by moving the stage with the sample along the beam axis closer or further away from the objective while recording optical microscope images at a z-distance of 1 µm. For each of these images a Laplacian filter is applied and the variance of the result is calculated, which serves as a measure of image sharpness. The stage is then moved to the z-position corresponding the maximum sharpness image and a fine-scan with 0.1 µm z-distance is performed in the range of +-1 µm in the same way as before to find the z-position of highest sharpness at a submicron precision. Since the focal planes of the optical microscope and the MALDI laser are aligned, this so found focal z-position is also the z-position where the MALDI-laser is in focus on the sample. This autofocus routine can be run either in brightfield mode, or in fluorescence mode. The python code of this program AutoFocusClass.py is provided in Data S1 in the supplementary information.

### In-source microscopy workflow

The scale factor to convert pixel on the camera sensor to position of the stage was calculated by using a 1951 USAF Resolution Test Target (Thorlabs). For selection of a region of interest (ROI) for an optical microscope scan in the ion source, autofocusing points are placed manually on the sample. The ROI is defined as the smallest enveloping rectangle that contains all points. The stage is then moved to each of the autofocusing points and an autofocus routine as described in “Autofocus of optics” is performed, which yields the focal z-offset of that x/y-coordinate. Using a radial basis function interpolation, the z-offset is calculated for every coordinate in a 10 µm x 10 µm mesh inside the ROI and saved to a z-interpolation file. After creating the z-interpolation, the optical microscope image of the ROI is taken. For this a 100 µm x 100 µm wide x/y-position mesh is created in the ROI. An optical microscope image is then taken at each mesh point at the z-offset of the closest point to the x/y-position from the z-interpolation file. To create a complete image of the ROI, an empty canvas of the correspondent size is created. Using the scale factor of the setup, images are cropped from the edges to a size of 100 µm x 100 µm and are placed onto the empty canvas using the center coordinates from the x/y-position mesh. After creating a z-interpolation file for a ROI once, multiple modalities, such as brightfield or different fluorescence channels can be recorded without creating a new interpolation. ImageJ was used to create overlay images of different modalities. The python code of this workflow can be found in the files RecordPositions.py, ZforRecordPositions.py, Scanning.py and stitching.py in Data S1 in the supplementary information.

### Transmission-MALDI-2 MSI

The MALDI-MSI run is setup partly using the stock flexImaging software (Bruker). Instead of using an external scan of the sample and teaching that to the stage coordinate system, the (multimodal) image of the ROI (see “In-source microscopy workflow”) is used. In the classic flexImaging workflow, three teaching points would be manually selected on the optical image. The corresponding stage coordinates would then be recorded by driving the sample-stage manually to the respective positions. Since center coordinates and pixel positions on the ROI image are already established from recording the ROI image, three x/y-positions from the 100 µm x 100 µm mesh (top left, bottom left, bottom right) and their corresponding stage coordinates are automatically deployed as teaching points. With these points, the setup file for the t-MALDI-2-MSI run can be automatically created, defeating any positioning errors introduced by manually selecting teaching points in the classic workflow. When the MALDI-laser is not directed at the exact middle pixel of the in-source optical microscope, fine adjustments of a few pixel offset can be applied at this stage. The automatically created setup file for the t-MALDI-2-MSI run can be opened directly in flexImaging to create measurement regions, select settings like pixel size, and laser shots per pixel. The previously acquired z-interpolation is recognized by timsControl software during t-MALDI-2-MSI measurements by providing the interpolation mesh as a geometry file. The python code for creating the flexImaging setup files can be found in Createmis.py Data S1 in the supplementary information. All t-MALDI-2-MSI experiments were performed with 25 laser shots per pixel and a pixel size of 1 x 1 µm^2^ for the cerebellum and cell culture samples and 2 x 2 µm^2^ for the tumor sample.

### Ion signal annotation

The lipid signals for the THP-1 cells were identified as described by Schwenzfeier et al. using shotgun lipidomics on a Q-Exactive Plus Orbitrap equipped with a dual ion funnel ^15,60^. Lipid extracts of 72 h PMA stimulated THP-1 macrophages were produced by an adapted protocol of Folch et al. ^61^. One million cells were centrifuged at 1000 g to separate them from the medium. Cells were resuspended in 1 ml CHCl_3_/MeOH (2:1 v/v) and lyzed for 10 min in an ultrasonic bath at room temperature followed by an incubation of 50 min at 4 °C in a cryoshaker at 500 RPM. Phase separation was induced by addition of 240 μl H_2_O and centrifugation at 9880 g for 10 min. The organic phase was transferred to a glas vial and the solvent was removed using a nitrogen evaporator. Lipid extracts were vacuum sealed and stored at -80 °C. For analysis the lipid extracts were dissolved in 500 μl MeOH and measured using electrospray ionisation (ESI) with a flow rate of 5 µl/min in positive and negative ion mode. A data dependent analysis workflow (DDA) was applied using tandem MS based on collision induced dissociation (CID) with stepped normalization collision energies (NCE) of 20, 25, 30 and an exclusion time of 300 s. The mass resolving power was set to 280 000 with an AGC target of 1e6 and a maximum injection time of 250 ms. The resulting data was merged with a t-MALDI-2-MSI dataset and imported into Lipostar MSI (Molecular Horizons) for annotation. The resulting feature list was hand-curated, which involved removing of isotope peaks and addition of known species and DHAP adducts that were not detectable in ESI.

Lipid signal annotation in mouse cerebellum was based on accurate mass (>3 ppm) and matched with literature data from full lipidomics analysis of mouse cerebellum ^30,33^. Because t-MALDI-2-MSI without the use of additional techniques like OzID or Paternò-Büchi-reactions ^62,63^ is not able to differentiate isomers within a lipid class (e.g. PC(16:0/18:1) and PC(16:1/18:0)) or between lipid classes (e.g. PE(16:0/18:0) and PC(15:0/16:0)) the literature data was reduced to 255 unique *m/z*-values according to the here employed level of specificity. For this, all lipid classes except DAGs and cholesterol were searched for as [M+H]^+^-ions. For DAG and cholesterol the more common [M-H_2_O+H]^+^-ion was used. 67 *m/z*-values with a S/N > 3 in the mean spectrum of the brain region with their respective highest signal intensity were thus annotated.

In the 4T1 murine tumor section annotation was also based on accurate mass (>3 ppm). For this signals with S/N > 3 were extracted from the mean spectrum and annotated using the LipidMaps “Bulk Structure Search” ^64^. Sphingolipids, glycerolipids, sterol lipids and glycerophospholipids were selected as possible lipid classes either as [M+H]^+^- or [M-H_2_O+H]^+^-ions. The resulting annotation list was manually curated to remove biologically unlikely annotation (e.g. lipids with odd side chains) or lipid classes that are usually not detected in positive ion-mode (e.g. glycerophosphoinositols). Using this method a total of 73 *m/z*-values were annotated.

Lipid nomenclature and shorthand notation was reported according to Liebisch ^65^.

### Single cell mass spectra from tissue

The external and in-source fluorescence images were co-registered using SimpleITK ^66^. For initialization a landmark registration was used. Based on the resulting translation matrix an affine image registration was performed using Mattes mutual information as metric and the gradient descent algorithm as optimizer ^67^. The number of bins for the mutual information was set to 50, the parameters for the optimizer were set as follows: learning rate (1), number of iterations (50), convergence minimum value (1E-6) and convergence window (20). The sampling strategy was set to random and 1 % of the pixels were sampled. The resulting transformation matrix was combined with the initial translation matrix to result in a global transformation matrix. Due to the inherent link between the t-MALDI-2-MSI measurement and the in-source fluorescence images the global transformation matrix could be used to resample the external fluorescence images to fit the exact region of the t-MALDI-2-MSI measurement. Cell masks were then created by using DeepCell Mesmer based on the DAPI channel (Hoechst 33342, nuclei) and the FITC channel (CellMask™ Green Actin Tracking Stain, actin). With FISCAS, mass spectra were calculated for each cell by compiling MSI pixels based on their co-localized cell masks. MSI pixels assigned to multiple cell masks were not used in further processing. Morphometric parameters were acquired using CellProfiler.

## Acknowledgements

The authors would like to thank Dagmar Mense (University of Münster) for support with cell culture, Mathis Richter, Alexander Fengler and Vincent Suerdieck for support with mouse tissue and immunohistochemistry (all University of Münster), Benedikt Geier (Stanford University) and Tanja Bien (Bruker Daltonics) for helpful discussions, Arne Fütterer, Andreas Höhne, Raik Milautzki, Henning Peise, and Olaf Roithner (all Bruker Daltonics) for hard- and software support, and Sarah Weischer and Thomas Zobel (Münster Imaging Network Cells in Motion Interfaculty Centre) for support with image analysis. This work was supported by grants from the German Research Foundation (DFG) (project 449437943 (Z1 Project of the TRR332) to OS and KD and project 326945247 to JS and KD).

## Notes

### Competing Interest Statement

Marcel Niehaus and Jens Hoehndorf are employees of Bruker Daltonics.

