## Supplementary Information for "Histology-Guided Single-Cell Mass Spectrometry Imaging using Integrated Bright-field and Fluorescence Microscopy"

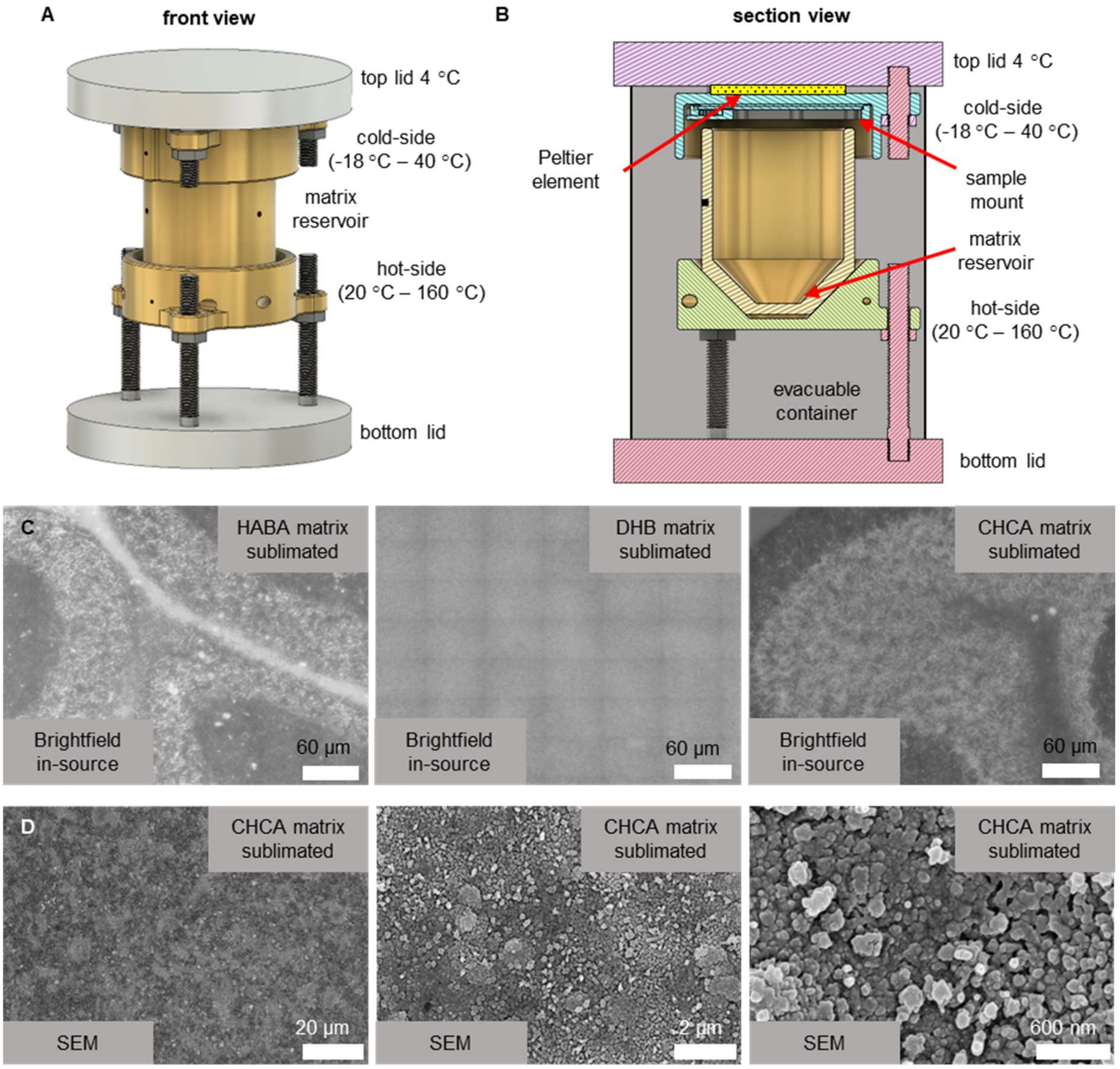


**Figure S1: Homogenous coating of samples with MALDI-matrix by resublimation.**
(A) Schematic of the custom-designed resublimation chamber. The top lid is actively water-cooled to 4 °C and serves as a heatsink for a Peltier element, which can heat up or chill the cold-side from -18 °C to 40 °C. The hot-side is heated by 3 heater cartridges with a total of 300 W heating power. Temperatures of the cold-side and hot-side are monitored using NTC thermistors. (B) Section view of the custom-designed resublimation chamber. This view of the resublimation chamber reveals the inner details of the setup. The sample can be magnetically clamped to the cold-side. The removable matrix reservoir is contacted to the hot-side using thermally conductive pads. An aluminum tube is mounted between the top and bottom lid to form an evacuable container. Using a turbomolecular pump, the whole setup can be evacuated to 10-2 mbar. (C) In-source BF image of three samples of mouse cerebellum coated with different MALDI matrices by resublimation. The choice of MALDI-matrix heavily influenced the quality of the in-source BF image. With 2-(4-Hydroxyphenylazo)benzoic acid (HABA) and CHCA matrix, the layers of the cerebellum can be easily defined, while resublimation with 2,5-Dihydroxybenzoic acid (DHB) prevents recording of a decent contrast image. (D) Scanning electron microscopy (SEM) images of a mouse cerebellum coated with CHCA MALDI matrix by resublimation. At three different levels of magnification, the homogenous matrix coating as well as the small matrix crystal size can be observed. A homogeneous coating is crucial to minimize local artifacts in t-MALDI-2 MSI. The matrix crystal size is between 50 nm and 200 nm and therefore is not likely to noticably limit the t-MALDI-2 MSI lateral resolution. SEM images were taken by a field emission SEM (Hitachi S4700) equipped with a digital scanning acquisition system (DISS5 point electronic)

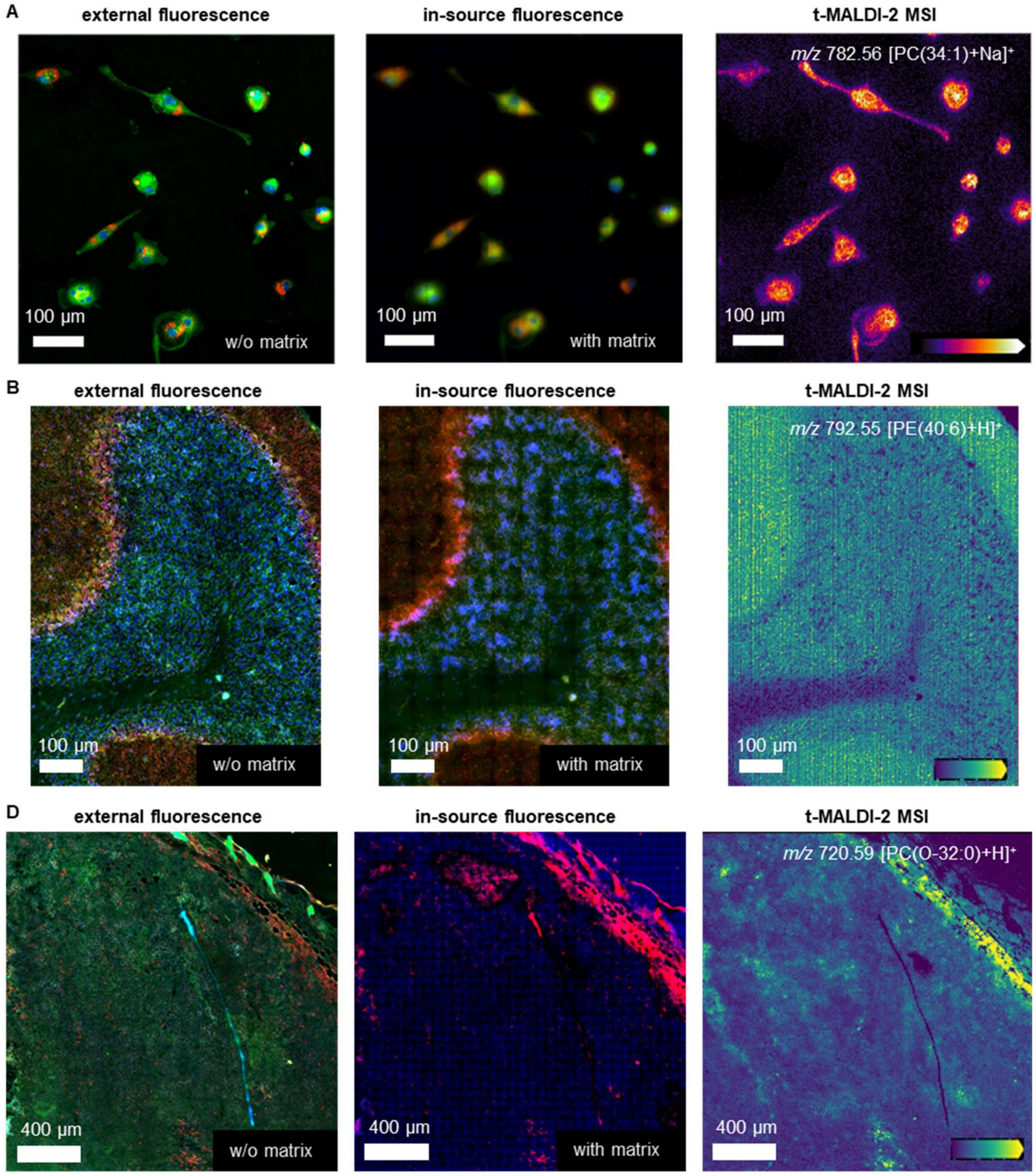


**Figure S2: Co-registration of external fluorescence image to t-MALDI-2 MSI across the different sample systems.**
(A) THP-1 derived macrophages. Initially, an external fluorescence image is obtained using three specific stains: Hoechst 33342 (nuclei, DAPI channel, blue), CellMask™ Green Actin Tracking Stain (actin, FITC channel, green), and LipidSpot™ 610 (lipid droplets, Cy3 channel, red). The sample is then coated with CHCA MALDI matrix via resublimation. Subsequently, the sample is placed into the ion source. A second fluorescence image, known as the in-source fluorescence image, is then taken using the same channels. The resulting t-MALDI-2 MSI is automatically aligned with the in-source fluorescence image due to the shared optics and sample stage, ensuring precise data alignment. During post-processing, the external fluorescence image is precisely co-registered to the in-source image and, consequently, to the t-MALDI-2 MSI data. (B) Mouse cerebellum: To begin, an external fluorescence image is acquired using the following stains: Hoechst 33342 (nuclei, DAPI channel, blue), CellMask™ Green Actin Tracking Stain (actin, FITC channel, green), and Alexa Fluor® 594 Anti-calbindin antibody (calbindin, Cy3 channel, red). The sample is subsequently coated with CHCA MALDI matrix by resublimation and loaded into the ion source, where an in-source fluorescence image is acquired using the same channels. The t-MALDI-2 MSI data is inherently co-registered with the in-source fluorescence image, facilitated by the consistent use of optics and the sample stage, ensuring accurate alignment of data sets. In post-processing, the external fluorescence image is effectively co-registered with the in-source fluorescence and the t-MALDI-2 MSI data. (C) Murine 4T1 tumor: An external fluorescence image is first captured using Hoechst 33342 (nuclei, DAPI channel, blue), CellMask™ Green Actin Tracking Stain (actin, FITC channel, green), and an immunofluorescence stain for neutrophil cells (Anti-Ly6G, Cy3 channel, red). The sample is then coated with CHCA MALDI matrix using the resublimation method and introduced into the ion source. Here, a second fluorescence image, termed the in-source fluorescence image, is recorded with the same fluorescence channels. The t-MALDI-2 MSI data is inherently aligned with this in-source fluorescence image due to the consistent optics and sample stage setup. In post-processing, precise co-registration of the external fluorescence image with the in-source fluorescence image, and therefore the t-MALDI-2 MSI data, is achieved.

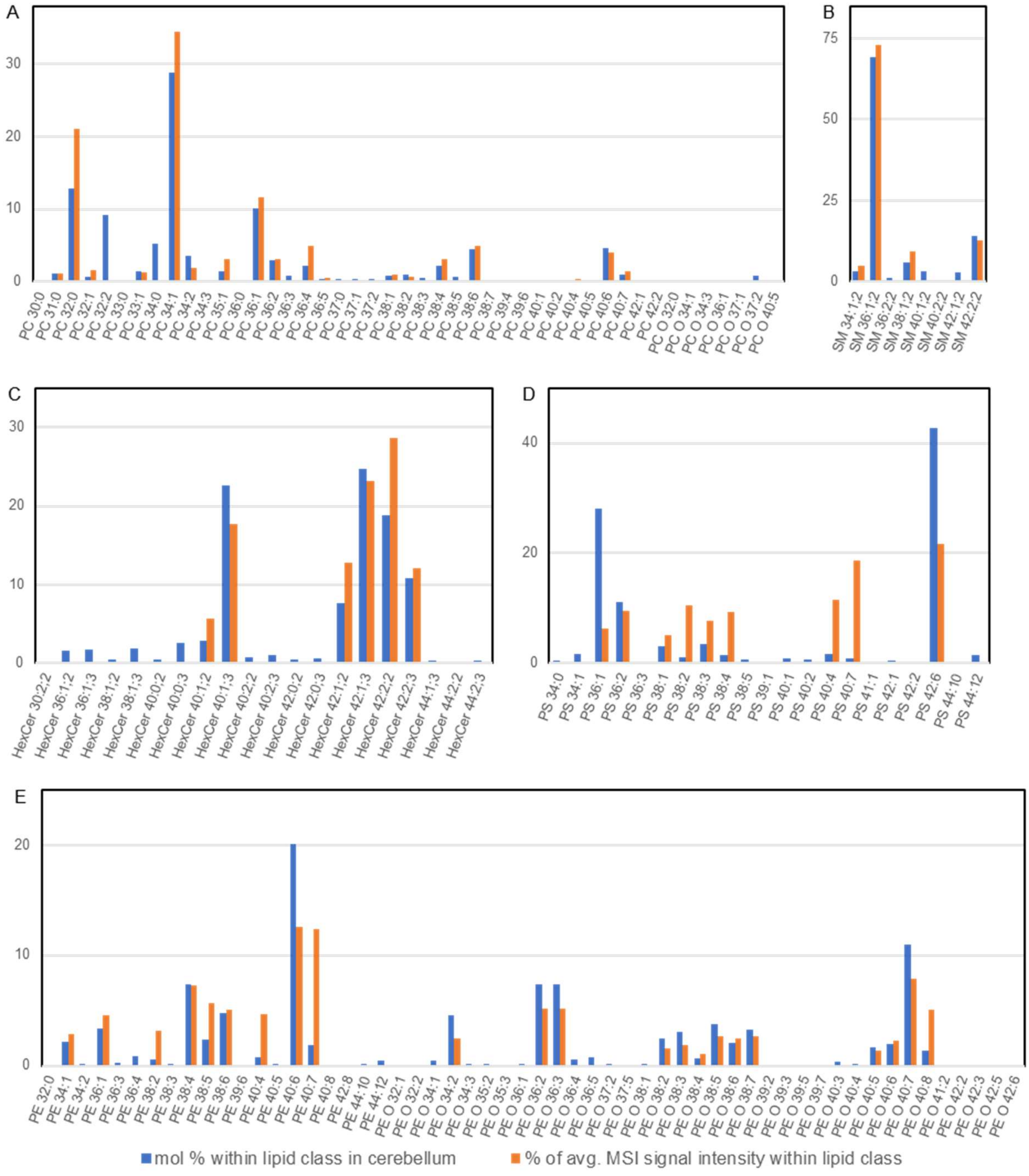


**Figure S3: Comparison of lipid signal intensities from murine cerebellum acquired with t-MALDI-MSI and values from the literature [44]**

(A) Molar percentage of different types of phosphatidylcholines (PC) in cerebellum derived from Fitzner *et al.*[44] in comparison with percentages of signal intensities for PCs detected in t-MALDI-2 MSI from the whole region depicted in figure 4. In both cases values are normalized to the sum of all reported/detected species from this lipid class.

(B) Comparison of molar percentage and percentage of signal intensities from t-MALDI-2-MSI for sphingomyelin (SM) species; analogous to (A)

(C) Comparison of molar percentage and percentage of signal intensities from t-MALDI-2-MSI for hexosylceramide (HexCer) species; analogous to (A)

(D) Comparison of molar percentage and percentage of signal intensities from t-MALDI-2-MSI for phosphatidylserine (PS) species; analogous to (A)

(E) Comparison of molar percentage and percentage of signal intensities from t-MALDI-2-MSI for phosphatidylethanolamine (PE) species; analogous to (A)

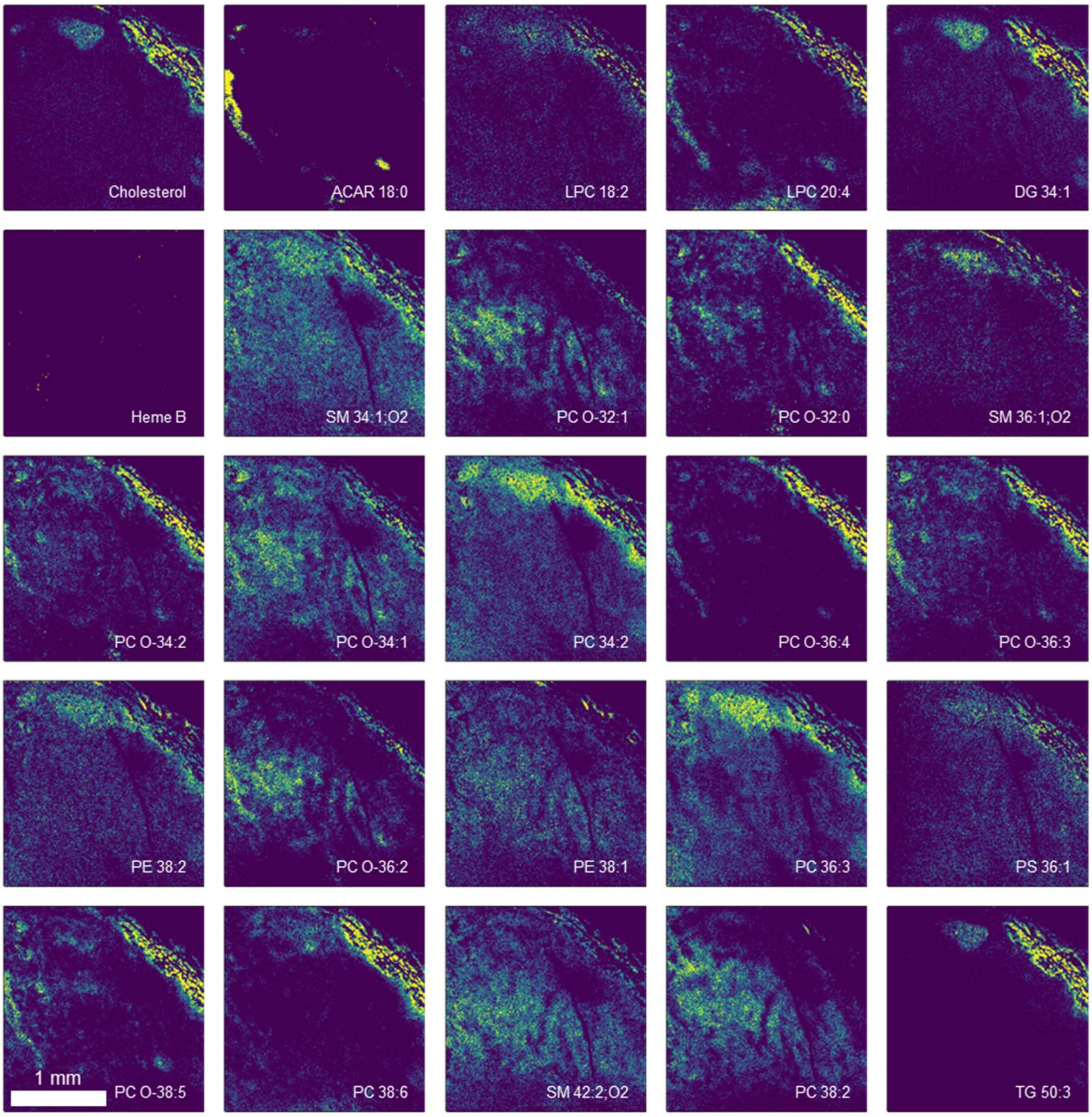


**Figure S4: Murine 4T1 tumor t-MALDI-2 MSI overview.**
Ion signal intensity distributions of different molecules from the measurement presented in Figure 5. Annotation was performed according to “ion signal annotation” found in the methods.

**Table S1:**
Tentative assignments of lipid signals in single cell analysis of THP-1 derived macrophages based on the comparison with results from shotgun lipidomics analysis of cellular extracts of the same cell type

| *m/z* exp. | *m/z* theo. | Delta | ppm | Name | Ion |
|---|---|---|---|---|---|
| **551.503** | 551.503 | 0.000 | 0.7 | DG 32:0 | [M+H-H2O]+ |
| **577.519** | 577.519 | 0.000 | 0.0 | DG 34:1 | [M+H-H2O]+ |
| **703.574** | 703.575 | 0.001 | 1.1 | SM 34:1;O2 | [M+H]+ |
| **723.494** | 723.496 | 0.002 | 2.6 | PA 38:5/PC 36:4-N(CH3)3 | [M+H]+ |
| **725.556** | 725.557 | 0.001 | 1.1 | SM 34:1;O2 | [M+Na]+ |
| **732.553** | 732.554 | 0.001 | 1.1 | PC 32:1 | [M+H]+ |
| **734.569** | 734.569 | 0.000 | 0.5 | PC 32:0 | [M+H]+ |
| **744.553** | 744.554 | 0.001 | 1.1 | PE 36:2 | [M+H]+ |
| **746.569** | 746.569 | 0.000 | 0.5 | PE 36:1 | [M+H]+ |
| **754.536** | 754.538 | 0.002 | 2.8 | PC 34:4 | [M+H]+ |
| **756.552** | 756.554 | 0.002 | 2.4 | PC 34:3 | [M+H]+ |
| **760.585** | 760.585 | 0.000 | 0.1 | PC 34:1 | [M+H]+ |
| **768.588** | 768.590 | 0.002 | 2.9 | PC O-36:4 | [M+H]+ |
| **770.605** | 770.606 | 0.001 | 1.7 | PC O-36:3 | [M+H]+ |
| **774.600** | 774.601 | 0.001 | 1.5 | PE 38:1 | [M+H]+ |
| **780.552** | 780.554 | 0.002 | 2.3 | PC 36:5 | [M+H]+ |
| **782.568** | 782.569 | 0.002 | 2.4 | PC 36:4 | [M+H]+ |
| **786.600** | 786.601 | 0.001 | 1.5 | PC 36:2 | [M+H]+ |
| **788.616** | 788.616 | 0.000 | 0.5 | PC 36:1 | [M+H]+ |
| **802.536** | 802.538 | 0.002 | 2.6 | PC 38:8 | [M+H]+ |
| **804.552** | 804.554 | 0.002 | 2.2 | PC 38:7 | [M+H]+ |
| **806.567** | 806.569 | 0.002 | 3.0 | PC 38:6 | [M+H]+ |
| **808.583** | 808.585 | 0.002 | 2.6 | PC 38:5 | [M+H]+ |
| **810.599** | 810.601 | 0.002 | 2.7 | PC 38:4 | [M+H]+ |
| **827.711** | 827.712 | 0.002 | 2.2 | TG 50:4 | [M+H]+ |
| **829.726** | 829.728 | 0.002 | 2.4 | TG 50:3 | [M+H]+ |
| **830.567** | 830.569 | 0.002 | 2.9 | PC 40:8 | [M+H]+ |
| **832.583** | 832.585 | 0.002 | 2.5 | PC 40:7 | [M+H]+ |
| **834.599** | 834.601 | 0.002 | 2.6 | PC 40:6 | [M+H]+ |
| **853.726** | 853.728 | 0.002 | 2.9 | TG 52:5 | [M+H]+ |
| **855.741** | 855.744 | 0.002 | 2.6 | TG 52:4 | [M+H]+ |
| **857.757** | 857.759 | 0.002 | 2.7 | TG 52:3 | [M+H]+ |
| **879.742** | 879.744 | 0.002 | 2.4 | TG 54:6 | [M+H]+ |
| **881.758** | 881.759 | 0.001 | 1.5 | TG 54:5 | [M+H]+ |
| **883.773** | 883.775 | 0.002 | 2.1 | TG 54:4 | [M+H]+ |
| **885.789** | 885.791 | 0.002 | 1.8 | TG 54:3 | [M+H]+ |
| **903.742** | 903.744 | 0.002 | 2.3 | TG 56:8 | [M+H]+ |
| **905.758** | 905.759 | 0.001 | 1.4 | TG 56:7 | [M+H]+ |
| **907.773** | 907.775 | 0.002 | 2.1 | TG 56:6 | [M+H]+ |
| **909.789** | 909.791 | 0.001 | 1.3 | TG 56:5 | [M+H]+ |
| **911.805** | 911.806 | 0.001 | 1.3 | TG 56:4 | [M+H]+ |

**Table S2:**
Tentative assignments of lipid signals in single cell analysis of murine cerebellum compared to intensity values from the literature; the percent in lipid class is calculated with respect to all lipid species in the respective lipid class based on the intensity provided in the literature or the single-cell analysis, respectively

| *m/z* exp. | *m/z* theo. | Delta | ppm | Name | Ion | intensity according to Fitzner et al. [30] | percent in lipid class [30] | Intensity from single-cell analysis | percent in lipid class |
|---|---|---|---|---|---|---|---|---|---|
| **369.352** | 369.352 | 0.000 | 1.1 | Cholesterol | [M+H-H2O]+ | 9517.0 | 100% | 697.1 | 100% |
| **496.341** | 496.340 | 0.001 | 2.6 | LPC 16:0 | [M+H]+ | 28.9 | 45% | 284.7 | 100% |
| **577.518** | 577.519 | 0.001 | 1.8 | DAG 34:1 | [M+H-H2O]+ | 44.3 | 17% | 504.3 | 55% |
| **599.503** | 599.503 | 0.000 | 0.6 | DAG 36:4 | [M+H-H2O]+ | 21.1 | 8% | 51.9 | 6% |
| **605.549** | 605.550 | 0.001 | 2.2 | DAG 36:1 | [M+H-H2O]+ | 36.6 | 14% | 225.5 | 25% |
| **627.534** | 627.535 | 0.001 | 1.1 | DAG 38:4 | [M+H-H2O]+ | 86.1 | 33% | 127.8 | 14% |
| **702.543** | 702.543 | 0.000 | 0.3 | PE O-34:2 | [M+H]+ | 318.9 | 5% | 59.7 | 2% |
| **703.575** | 703.575 | 0.000 | 0.3 | SM 34:1;2 | [M+H]+ | 19.2 | 3% | 26.2 | 5% |
| **718.538** | 718.538 | 0.000 | 0.1 | PE 34:1 | [M+H]+ | 152.1 | 2% | 69.8 | 3% |
| **720.554** | 720.554 | 0.000 | 0.4 | PC 31:0 | [M+H]+ | 126.6 | 1% | 144.3 | 1% |
| **728.558** | 728.559 | 0.001 | 1.1 | PE O-36:3 | [M+H]+ | 517.0 | 7% | 124.2 | 5% |
| **730.538** | 730.538 | 0.000 | 0.1 | PC 32:2 | [M+H]+ | 999.1 | 9% | 31.3 | 0% |
| **730.574** | 730.574 | 0.000 | 0.7 | PE O-36:2 | [M+H]+ | 519.8 | 7% | 125.3 | 5% |
| **731.606** | 731.606 | 0.000 | 0.2 | SM 36:1;2 | [M+H]+ | 417.8 | 69% | 394.3 | 73% |
| **732.554** | 732.554 | 0.000 | 0.4 | PC 32:1 | [M+H]+ | 70.3 | 1% | 198.6 | 2% |
| **734.570** | 734.569 | 0.001 | 0.8 | PC 32:0 | [M+H]+ | 1398.1 | 13% | 2664.4 | 21% |
| **744.554** | 744.554 | 0.000 | 0.4 | PE 36:1 | [M+H]+ | 234.4 | 3% | 109.6 | 5% |
| **746.569** | 746.569 | 0.000 | 0.5 | PC 33:1 | [M+H]+ | 161.3 | 1% | 156.1 | 1% |
| **748.526** | 748.528 | 0.002 | 2.0 | PE O-38:7 | [M+H]+ | 227.0 | 3% | 64.4 | 3% |
| **749.533** | 749.533 | 0.000 | 0.5 | PG 34:1 | [M+H]+ | 22.3 | 48% | 58.2 | 100% |
| **750.541** | 750.543 | 0.002 | 2.9 | PE O-38:6 | [M+H]+ | 147.3 | 2% | 58.1 | 2% |
| **752.559** | 752.559 | 0.000 | 0.2 | PE O-38:5 | [M+H]+ | 263.8 | 4% | 63.7 | 3% |
| **754.573** | 754.574 | 0.001 | 2.0 | PE O-38:4 | [M+H]+ | 49.7 | 1% | 24.6 | 1% |
| **756.591** | 756.590 | 0.001 | 1.2 | PE O-38:3 | [M+H]+ | 214.9 | 3% | 44.2 | 2% |
| **758.569** | 758.569 | 0.000 | 0.5 | PC 34:2 | [M+H]+ | 386.7 | 4% | 231.4 | 2% |

| | | | | | | | | | |
|---|---|---|---|---|---|---|---|---|---|
| **758.606** | 758.606 | 0.000 | 0.3 | PE O-38:2 | [M+H]+ | 175.1 | 2% | 38.1 | 2% |
| **759.638** | 759.637 | 0.001 | 0.8 | SM 38:1;2 | [M+H]+ | 35.9 | 6% | 51.0 | 9% |
| **760.585** | 760.585 | 0.000 | 0.0 | PC 34:1 | [M+H]+ | 3128.4 | 29% | 4344.1 | 34% |
| **764.521** | 764.522 | 0.001 | 1.9 | PE 38:6 | [M+H]+ | 333.2 | 5% | 121.8 | 5% |
| **766.536** | 766.538 | 0.002 | 2.7 | PE 38:5 | [M+H]+ | 163.5 | 2% | 135.5 | 6% |
| **768.554** | 768.554 | 0.000 | 0.3 | PE 38:4 | [M+H]+ | 518.3 | 7% | 175.7 | 7% |
| **772.584** | 772.585 | 0.001 | 1.3 | PE 38:2 | [M+H]+ | 37.2 | 1% | 77.2 | 3% |
| **774.542** | 774.543 | 0.001 | 1.5 | PE O-40:8 | [M+H]+ | 98.9 | 1% | 121.4 | 5% |
| **774.601** | 774.601 | 0.000 | 0.4 | PC 35:1 | [M+H]+ | 159.2 | 1% | 384.9 | 3% |
| **776.558** | 776.559 | 0.001 | 1.1 | PE O-40:7 | [M+H]+ | 772.9 | 11% | 188.5 | 8% |
| **778.574** | 778.574 | 0.000 | 0.6 | PE O-40:6 | [M+H]+ | 139.5 | 2% | 53.8 | 2% |
| **780.553** | 780.554 | 0.001 | 0.9 | PC 36:5 | [M+H]+ | 38.6 | 0% | 69.1 | 1% |
| **780.589** | 780.590 | 0.001 | 1.4 | PE O-40:5 | [M+H]+ | 118.4 | 2% | 31.8 | 1% |
| **782.569** | 782.569 | 0.000 | 0.5 | PC 36:4 | [M+H]+ | 232.3 | 2% | 615.9 | 5% |
| **784.664** | 784.666 | 0.002 | 2.6 | HexCer 40:1;2 | [M+H]+ | 33.8 | 3% | 22.4 | 6% |
| **786.601** | 786.601 | 0.000 | 0.4 | PC 36:2 | [M+H]+ | 322.8 | 3% | 380.8 | 3% |
| **788.544** | 788.544 | 0.000 | 0.6 | PS 36:2 | [M+H]+ | 448.1 | 11% | 54.4 | 9% |
| **788.617** | 788.616 | 0.001 | 0.8 | PC 36:1 | [M+H]+ | 1102.6 | 10% | 1466.6 | 12% |
| **790.537** | 790.538 | 0.001 | 1.4 | PE 40:7 | [M+H]+ | 131.1 | 2% | 298.2 | 12% |
| **790.559** | 790.559 | 0.000 | 0.3 | PS 36:1 | [M+H]+ | 1144.7 | 28% | 35.7 | 6% |
| **792.554** | 792.554 | 0.000 | 0.3 | PE 40:6 | [M+H]+ | 1413.0 | 20% | 302.6 | 13% |
| **796.585** | 796.585 | 0.000 | 0.0 | PE 40:4 | [M+H]+ | 54.9 | 1% | 111.4 | 5% |
| **800.661** | 800.661 | 0.000 | 0.0 | HexCer 40:1;3 | [M+H]+ | 264.4 | 23% | 70.2 | 18% |
| **806.570** | 806.569 | 0.001 | 0.8 | PC 38:6 | [M+H]+ | 477.4 | 4% | 622.1 | 5% |
| **810.601** | 810.601 | 0.000 | 0.4 | PC 38:4 | [M+H]+ | 235.6 | 2% | 382.5 | 3% |
| **810.682** | 810.682 | 0.000 | 0.4 | HexCer 42:2;2 | [M+H]+ | 220.5 | 19% | 113.2 | 29% |
| **812.543** | 812.544 | 0.001 | 0.7 | PS 38:4 | [M+H]+ | 56.7 | 1% | 54.0 | 9% |
| **812.695** | 812.697 | 0.002 | 2.9 | HexCer 42:1;2 | [M+H]+ | 89.7 | 8% | 50.7 | 13% |
| **813.687** | 813.684 | 0.003 | 3.2 | SM 42:2;2 | [M+H]+ | 84.3 | 14% | 69.4 | 13% |

| | | | | | | | | | |
|---|---|---|---|---|---|---|---|---|---|
| **814.559** | 814.559 | 0.000 | 0.3 | PS 38:3 | [M+H]+ | 143.6 | 4% | 44.4 | 8% |
| **814.632** | 814.632 | 0.000 | 0.0 | PC 38:2 | [M+H]+ | 101.5 | 1% | 77.9 | 1% |
| **816.575** | 816.575 | 0.000 | 0.2 | PS 38:2 | [M+H]+ | 42.5 | 1% | 60.4 | 10% |
| **816.648** | 816.648 | 0.000 | 0.4 | PC 38:1 | [M+H]+ | 82.0 | 1% | 116.7 | 1% |
| **818.590** | 818.591 | 0.001 | 0.6 | PS 38:1 | [M+H]+ | 123.0 | 3% | 28.8 | 5% |
| **826.677** | 826.677 | 0.000 | 0.5 | HexCer 42:2;3 | [M+H]+ | 126.4 | 11% | 47.7 | 12% |
| **828.693** | 828.692 | 0.001 | 0.9 | HexCer 42:1;3 | [M+H]+ | 289.2 | 25% | 91.6 | 23% |
| **832.585** | 832.585 | 0.000 | 0.0 | PC 40:7 | [M+H]+ | 110.5 | 1% | 181.2 | 1% |
| **834.526** | 834.528 | 0.002 | 2.3 | PS 40:7 | [M+H]+ | 31.6 | 1% | 107.9 | 19% |
| **834.601** | 834.601 | 0.000 | 0.4 | PC 40:6 | [M+H]+ | 493.7 | 5% | 514.0 | 4% |
| **836.543** | 836.544 | 0.001 | 0.7 | PS 42:6 | [M+H]+ | 1741.7 | 43% | 125.8 | 22% |
| **838.631** | 838.632 | 0.001 | 1.2 | PC 40:4 | [M+H]+ | 23.4 | 0% | 40.2 | 0% |
| **840.573** | 840.575 | 0.002 | 2.2 | PS 40:4 | [M+H]+ | 67.5 | 2% | 66.3 | 11% |

**Table S1:**
Tentative assignments of lipid signals of the investigated 4T1 tumor sample

| *m/z* exp. | *m/z* theo. | Delta | ppm | Name | Ion |
|---|---|---|---|---|---|
| **369.352** | 369.352 | 0.000 | 0.3 | Cholesterol | [M+H-H2O]+ |
| **428.374** | 428.373 | 0.001 | 1.4 | ACAR 18:0 | [M+H]+ |
| **494.324** | 494.324 | 0.001 | 1.2 | LPC 16:1 | [M+H]+ |
| **496.340** | 496.340 | 0.000 | 0.6 | LPC 16:0 | [M+H]+ |
| **520.340** | 520.340 | 0.000 | 0.6 | LPC 18:2 | [M+H]+ |
| **522.355** | 522.355 | 0.000 | 0.8 | LPC 18:1 | [M+H]+ |
| **524.371** | 524.371 | 0.001 | 1.1 | LPC 18:0 | [M+H]+ |
| **544.339** | 544.340 | 0.001 | 1.5 | LPC 20:4 | [M+H]+ |
| **549.487** | 549.488 | 0.001 | 2.2 | DG 32:1 | [M+H-H2O]+ |
| **551.503** | 551.503 | 0.001 | 1.6 | DG 32:0 | [M+H-H2O]+ |
| **575.502** | 575.503 | 0.001 | 2.4 | DG 34:2 | [M+H-H2O]+ |
| **577.518** | 577.519 | 0.001 | 1.7 | DG 34:1 | [M+H-H2O]+ |
| **597.487** | 597.488 | 0.001 | 2.0 | DG 36:5 | [M+H-H2O]+ |
| **599.502** | 599.503 | 0.001 | 2.3 | DG 36:4 | [M+H-H2O]+ |
| **601.518** | 601.519 | 0.001 | 1.7 | DG 36:3 | [M+H-H2O]+ |
| **603.533** | 603.535 | 0.002 | 2.8 | DG 36:2 | [M+H-H2O]+ |
| **605.549** | 605.550 | 0.002 | 3.0 | DG 36:1 | [M+H-H2O]+ |
| **625.518** | 625.519 | 0.001 | 2.4 | DG 38:5 | [M+H-H2O]+ |
| **627.533** | 627.535 | 0.002 | 2.7 | DG 38:4 | [M+H-H2O]+ |
| **701.559** | 701.559 | 0.000 | 0.3 | SM 34:2;O2 | [M+H]+ |
| **703.575** | 703.575 | 0.000 | 0.4 | SM 34:1;O2 | [M+H]+ |
| **704.522** | 704.523 | 0.000 | 0.7 | PC 30:1 | [M+H]+ |
| **706.538** | 706.538 | 0.000 | 0.1 | PC 30:0 | [M+H]+ |
| **718.538** | 718.538 | 0.001 | 0.8 | PE 34:1 | [M+H]+ |
| **718.575** | 718.575 | 0.001 | 0.7 | PC O-32:1 | [M+H]+ |
| **720.554** | 720.554 | 0.000 | 0.4 | PE 34:0 | [M+H]+ |
| **720.589** | 720.590 | 0.001 | 1.7 | PC O-32:0 | [M+H]+ |
| **728.522** | 728.523 | 0.001 | 1.4 | PC 32:3 | [M+H]+ |
| **730.538** | 730.538 | 0.000 | 0.1 | PC 32:2 | [M+H]+ |
| **731.606** | 731.606 | 0.000 | 0.1 | SM 36:1;O2 | [M+H]+ |
| **732.554** | 732.554 | 0.000 | 0.3 | PC 32:1 | [M+H]+ |
| **734.569** | 734.569 | 0.000 | 0.5 | PC 32:0 | [M+H]+ |
| **742.538** | 742.538 | 0.001 | 0.8 | PE 36:3 | [M+H]+ |
| **744.553** | 744.554 | 0.001 | 1.1 | PE 36:2 | [M+H]+ |
| **744.590** | 744.590 | 0.000 | 0.3 | PC O-34:2 | [M+H]+ |
| **746.569** | 746.569 | 0.000 | 0.5 | PE 36:1 | [M+H]+ |
| **746.606** | 746.606 | 0.000 | 0.3 | PC O-34:1 | [M+H]+ |
| **748.584** | 748.585 | 0.001 | 1.5 | PE 36:0 | [M+H]+ |
| **754.537** | 754.538 | 0.001 | 1.5 | PC 34:4 | [M+H]+ |
| **756.554** | 756.554 | 0.000 | 0.3 | PC 34:3 | [M+H]+ |
| **758.570** | 758.569 | 0.001 | 0.8 | PC 34:2 | [M+H]+ |
| **760.585** | 760.585 | 0.000 | 0.1 | PC 34:1 | [M+H]+ |
| **768.553** | 768.554 | 0.001 | 1.0 | PE 38:4 | [M+H]+ |

| | | | | | |
|---|---|---|---|---|---|
| **768.590** | 768.590 | 0.000 | 0.3 | PC O-36:4 | [M+H]+ |
| **770.569** | 770.569 | 0.001 | 1.2 | PE 38:3 | [M+H]+ |
| **770.605** | 770.606 | 0.001 | 1.7 | PC O-36:3 | [M+H]+ |
| **772.585** | 772.585 | 0.000 | 0.1 | PE 38:2 | [M+H]+ |
| **772.621** | 772.622 | 0.000 | 0.6 | PC O-36:2 | [M+H]+ |
| **774.601** | 774.601 | 0.000 | 0.4 | PE 38:1 | [M+H]+ |
| **780.554** | 780.554 | 0.000 | 0.4 | PC 36:5 | [M+H]+ |
| **782.570** | 782.569 | 0.000 | 0.1 | PC 36:4 | [M+H]+ |
| **784.585** | 784.585 | 0.000 | 0.1 | PC 36:3 | [M+H]+ |
| **786.601** | 786.601 | 0.000 | 0.4 | PC 36:2 | [M+H]+ |
| **788.616** | 788.616 | 0.001 | 1.1 | PC 36:1 | [M+H]+ |
| **790.560** | 790.559 | 0.001 | 0.9 | PS 36:1 | [M+H]+ |
| **794.607** | 794.606 | 0.001 | 0.9 | PC O-38:5 | [M+H]+ |
| **796.622** | 796.622 | 0.000 | 0.0 | PC O-38:4 | [M+H]+ |
| **798.635** | 798.637 | 0.002 | 2.6 | PC O-38:3 | [M+H]+ |
| **800.617** | 800.616 | 0.001 | 0.7 | PE 40:2 | [M+H]+ |
| **802.631** | 802.632 | 0.001 | 1.2 | PE 40:1 | [M+H]+ |
| **806.570** | 806.569 | 0.000 | 0.1 | PC 38:6 | [M+H]+ |
| **808.586** | 808.585 | 0.000 | 0.5 | PC 38:5 | [M+H]+ |
| **810.602** | 810.601 | 0.001 | 1.0 | PC 38:4 | [M+H]+ |
| **812.615** | 812.616 | 0.001 | 1.7 | PC 38:3 | [M+H]+ |
| **813.685** | 813.684 | 0.001 | 0.7 | SM 42:2;O2 | [M+H]+ |
| **814.632** | 814.632 | 0.000 | 0.6 | PC 38:2 | [M+H]+ |
| **816.646** | 816.648 | 0.002 | 2.1 | PC 38:1 | [M+H]+ |
| **827.713** | 827.712 | 0.000 | 0.2 | TG 50:4 | [M+H]+ |
| **829.728** | 829.728 | 0.000 | 0.0 | TG 50:3 | [M+H]+ |
| **834.601** | 834.601 | 0.000 | 0.4 | PC 40:6 | [M+H]+ |
| **855.746** | 855.744 | 0.002 | 2.2 | TG 52:4 | [M+H]+ |
| **879.745** | 879.744 | 0.001 | 1.6 | TG 54:6 | [M+H]+ |
| **881.760** | 881.759 | 0.001 | 0.8 | TG 54:5 | [M+H]+ |
